# Modelling temporal dynamics of genetic diversity in stage-structured plant populations with reference to demographic genetic structure

**DOI:** 10.1101/2021.11.30.470535

**Authors:** Yoichi Tsuzuki, Takenori Takada, Masashi Ohara

## Abstract

1. Predicting temporal dynamics of genetic diversity is important for assessing long-term population persistence. In stage-structured populations, especially in perennial plant species, genetic diversity is often compared among life history stages, such as seedlings, juveniles, and flowerings, using neutral genetic markers. Because individuals in mature stages will die and be replaced by those in more immature stages over the course of time, the comparison among stages (sometimes referred to as demographic genetic structure) has been regarded as a proxy of potential genetic changes that accompany the turnover of constituent individuals. However, because demographic genetic structure had not been theoretically examined, the basic property and the validity of demographic genetic structure remained unclear.
2. We developed a matrix model which was made up of difference equations of expected heterozygosity, a common proxy of genetic diversity, of each life history stage at a neutral locus in stage-structured plant populations. Based on the model, we formulated demographic genetic structure as well as the annual change rate of expected heterozygosity (denoted as *η*). We obtained theoretical expectation of demographic genetic structure and *η* from our model and compared them with computational results of stochastic simulation for randomly generated 3,000 life histories for model validation. We then examined the relationships of demographic genetic structure with effective population size *N_e_*, which is the determinants of diversity loss per generation time, as well as with *η*.
3. Theoretical expectations on *η* and demographic genetic structure fitted well to the results of stochastic simulation, supporting the validity of our model. Demographic genetic structure varied independently of *N_e_* and *η*, while having a strong correlation with stable stage distribution: expected heterozygosity was lower in stages with fewer individuals.
4. Our results indicate that demographic genetic structure strongly reflects stable stage distribution, rather than temporal genetic dynamics, and that inferring future genetic diversity solely from demographic genetic structure would be misleading. Instead of demographic genetic structure, the newly-defined statistics *η* will be an useful tool to predict genetic diversity at the same time scale as population dynamics, facilitating evaluation on population viability from a genetic point of view.

## 1 Introduction

Genetic diversity, or standing genetic variation, is a source of adaptive evolution (Barrett & Schluter, 2008). Populations with high genetic diversity are more likely to adapt to environmental changes and to persist for a long period (Agashe et al., 2011; Ramsayer et al., 2013). Therefore, it is necessary to examine the temporal dynamics of genetic diversity for assessing long-term population viability (Mimura et al., 2017).

The rate of change in genetic diversity per generation time is primarily determined by effective population size (*N_e_*): the larger *N_e_* is, the weaker genetic drift works, and more likely genetic diversity is maintained (Crow & Kimura, 1970). Although N_e_ is first theoretically proposed for populations without generation overlap, many wild populations including perennial plants have generation overlap and are made up of individuals differing in age or life history stage. Previous theoretical studies have applied the concept of effective population size to populations structured by age (Felsenstein, 1971; Hill, 1972; Hill, 1979; Johnson, 1977) or by stage (Orive, 1993; Yonezawa et al., 2000) by formulating *N_e_* with demographic rates (age- or stage-specific survival rates and fecundities). These formulations enable us to calculate Ne and to assess the temporal genetic dynamics in species with complex life histories (Waples et al., 2011; Waples et al., 2013).

Meanwhile, some empirical genetic studies haves not examined Ne directly to predict future genetic diversity. Instead, genetic diversity is comparatively estimated for each age or stage class at a single time point with neutral genetic markers (Aldrich et al., 1998; Ally & Ritland, 2007; Kettle et al., 2007; Linhart et al., 1981; Murren, 2003; Schmidt et al., 2018; Vranckx et al., 2014). The resultant within-population genetic structure (sometimes referred to as demographic genetic structure) is considered to reflect potential genetic changes that accompany the turnover of constituent individuals. For example, if juvenile stage is less diverse than more mature stages, genetic diversity would decrease with the replacement of mature individuals to juveniles. Because species with age or stage structure are mostly long-lived and long-term genetic monitoring is impractical, demographic genetic structure has been considered as a rough but a convenient empirical approach to infer the temporal genetic dynamics (Mimura et al., 2017; Schmidt et al., 2018).

Despite its empirical usage, mathematical and theoretical basis of demographic genetic structure has been in its infancy. Unlike N_e_, how demographic genetic structure is determined by demographic rates has not been formulated. Relationships with N_e_ have also remained unexplored, which raises a question on whether Ne and demographic genetic structure are largely redundant or highlight different aspects of temporal genetic dynamics. Moreover, lack of theoretical background draws concerns about the current interpretation on demographic genetic structure in terms of life history features of stage-structured populations. While analysis on demographic genetic structure implicitly assumes that individuals sequentially grow and die from juvenile to mature stage classes, this assumption is potentially invalid especially in perennial plants. In most perennial plant species, whose life histories are structured by stage, not by age (Silvertown, 1987), aging (or passing of time) does not necessarily promote growth and maturation. Some individuals might keep proceeding to more mature stages, while others remain in the same stage for a long period (stasis) or even reverse to more juvenile stages (retrogression). For example, long-lived woodland perennial herbs of the genus *Trillium* show stasis for more than ten years in juvenile stages as well as go back from a mature reproductive stage to a pre-reproductive one in response to resource exhaustion (Knight, 2004; Ohara et al., 2001; Tomimatsu & Ohara, 2010). The static and bidirectional flows in the life cycle complicate the order of individual turnover in a population. It has not been theoretically confirmed if demographic genetic structure still serves as a proxy for temporal changes despite these challenges. Mathematical formulation that encompasses demographic genetic structure, as well as the temporal change in genetic diversity, will provide integrative understandings on all the problems mentioned above, but has never been achieved so far.

In this study, we develop a matrix model to describe the temporal dynamics of expected heterozygosity, which is used as a common metric of genetic diversity, for a neutral locus of a stage-structured perennial plant species. The model is made up of difference equations of the probability that two genes randomly sampled from a given life history stage are non-identical-by-descent. Based on the model, we formulate demographic genetic structure and *N_e_*. Thus, our model allows integrative analysis on demographic genetic structure, temporal dynamics of genetic diversity, and their relationships. In the following sections, we describe the derivation procedures of our model (section 2.1), the validation of our model (section 2.2), and the assessment on whether demographic genetic structure serves as a good proxy for the temporal changes in genetic diversity (section 2.3).

## 2 Materials and Methods

### 2.1 Model development

#### 2.1.1 Overview

Felsenstein (1971) derived inbreeding effective population size for an age-structured population, which is also described in Charlesworth (2008). We partly follow mathematical formulation procedures in Felsenstein (1971) and Charlesworth (2008) to develop difference equations that describes the dynamics of the probability of non-identical-by-descent at a neutral locus for a closed, stage-structured population, supposing a diploid perennial plant species. We do not consider sex differences because most plants are hermaphrodite (Torices et al., 2011). Census population size is set to *N*, which is divided into *n* life history stages (*N_x_, N*_2_,…, *N_n_*).

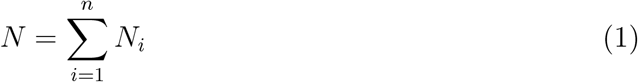

As in Felsenstein (1971), we assume an equilibrium state, where the census population size and its allocation to each stage (stage distribution) are constant over time. The probability of transition (either growth, stasis, or retrogression) from stage *j* to stage *i* is *t_ij_* per year. In each year, individuals randomly mate and *f_ij_* newborns join stage *i* from a parent in stage *j*. *a_ij_*, which denotes the sum of *t_ij_* and *f_ij_*, describes the total flow of individuals from stage *j* to *i* between successive years.

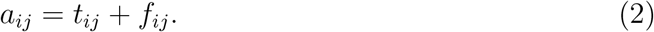

Population dynamics can be modeled by the following transition matrix.

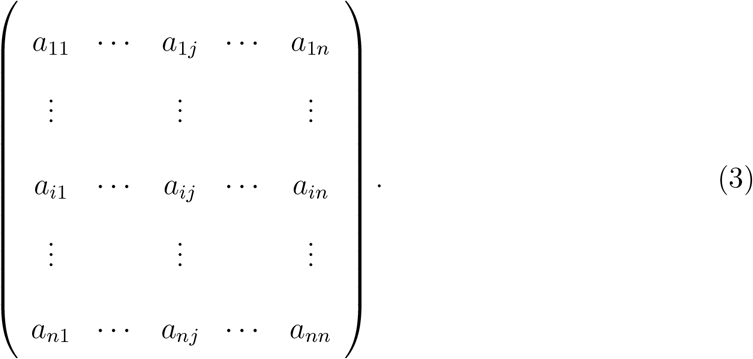

Stable stage distribution, which is the relative number of individuals among stages in the equilibrium state, is proportional to the leading right eigenvector of the transition matrix (Caswell, 2001).

We define *H_ij,t_* as the probability that two genes randomly sampled from stage *i* and *j* with replacement in year *t* are not identical-by-descent. *H_ij,t_* is equivalent to expected heterozygosity, which is a common proxy of genetic diversity in empirical studies. The goal of our model development is to formulate *H_ij,t_* for all possible *i* and *j*, which enables us to obtain theoretical counterpart of demographic genetic structure, that is, neutral genetic diversity of each stage class at a particular time point. Here, we provide only key equations in the main text for tidiness, while the complete derivation procedures are given in Supporting Information.

#### 2.1.2 Difference equations of *H_ij,t_*

We separately consider two mutually exclusive situations: *i* ≠ *j* (case 1) and *i* = *j* (case 2). Both cases can be further split into six situations. Firstly, two genes randomly sampled in year t were either in the same stage (say, stage *m,* case A) or in different stages (say, stage *k* and *l*, case B) in year *t* — 1. Furthermore, genes can move among stages either by survival (grow, stasis, and retrogression) or by reproduction. Survival and reproduction are essentially different because reproduction allows one gene to be replicated and to move to multiple stages simultaneously and independently, while survival does not. There are three possibilities in how the two genes sampled were transferred: both genes were transferred by survival (case *α*), one by survival and the other by reproduction (case *β*), and both by reproduction (case *γ*). Considering the combinations of where (case A and B) and how (case *α, β*, and *γ*) the two genes sampled came from, there are 6 mutually exclusive situations to be considered in both case 1 and 2 (Figure 1).

**Figure 1:**
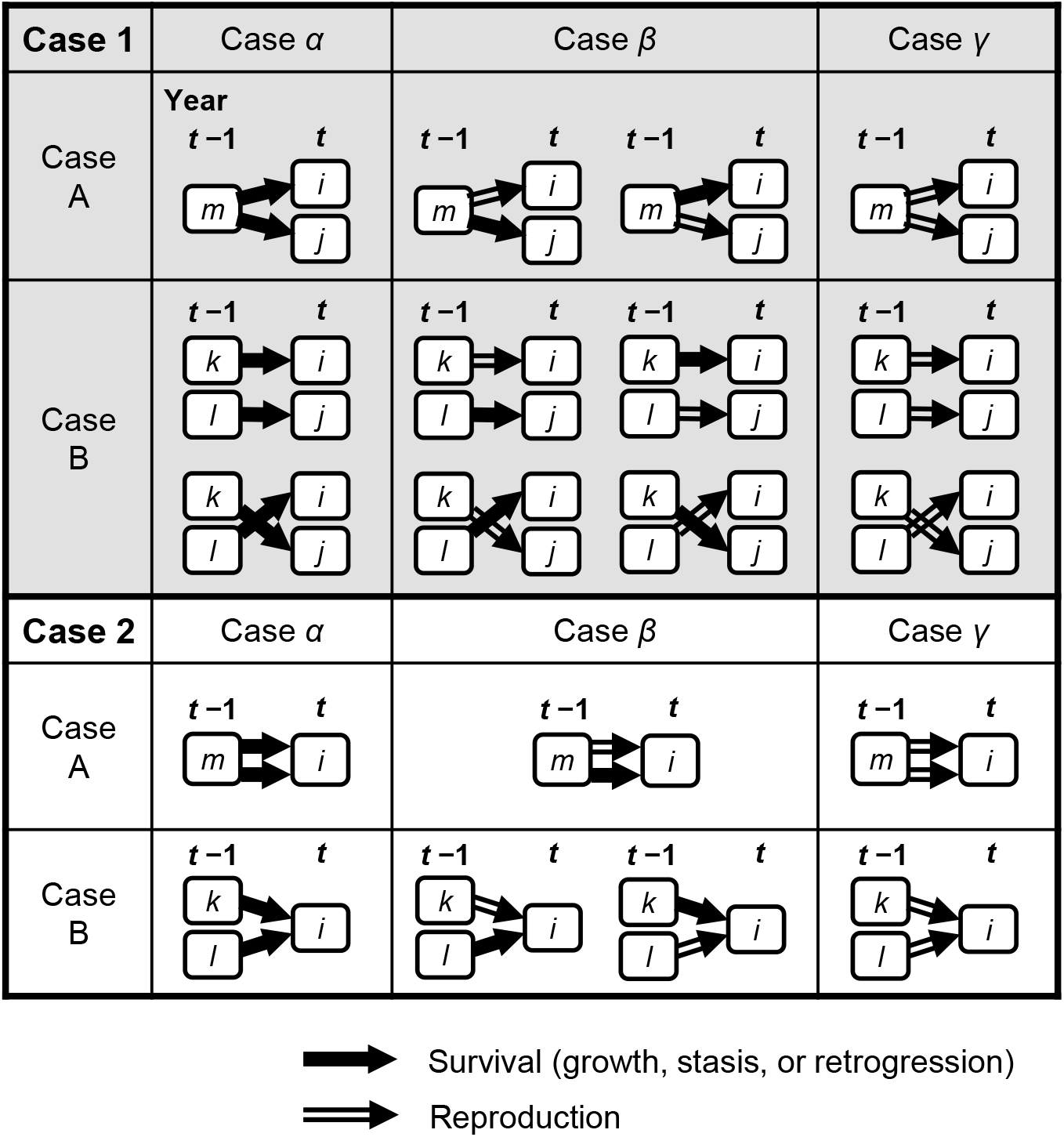
Temporal trajectories from time *t* — 1 to *t* with regard to the two genes sampled in time t. Rounded rectangles stand for life history stages. Arrows stand for the temporal movements of genes either by survival (single line) or reproduction (double line). There are 12 mutually exclusive situations based on three criteria: (1) whether the destinations are different (case 1, shown on gray background) or not (case 2, shown on white); (2) whether the origins are the same (case A) or not (case B); (3) how the two genes were transferred (case α: survival; case *β*: survival and reproduction; case γ: reproduction)

In case 1, *H_ij,t_* can be decomposed as follows.

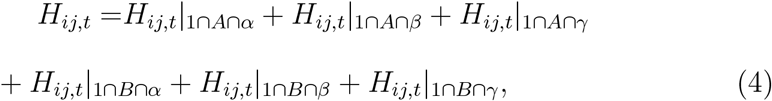

where the cap symbol ⋂ stands for the co-occurrence of multiple cases: 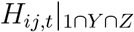 stands for *H_ij,t_* that simultaneously satisfies case 1, Y, and Z (*Y* = *A,B*; *Z* = *α,β,γ*). All six 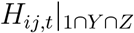 on the right side of equation 4 are formulated as follows (see Supporting Information 1.1).

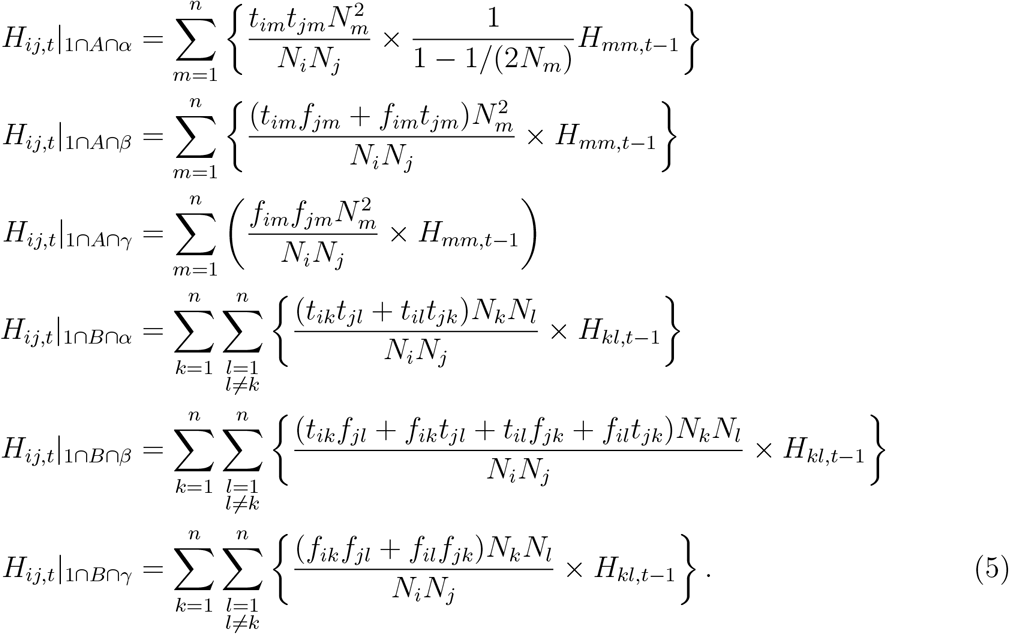

Each 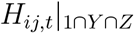 is shown as a summation of a multiplications of two terms. The first term is a conditional probability of case 1 ⋂ *Y* ⋂ *Z* given case 1. For example, the first term of 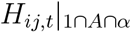 can be rewritten as 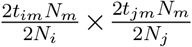, which is the number of two-gene pairs that fall into case 1, A, and *α* simultaneously under a specific *m* (2*t_im_N_m_* × 2*t_jm_N_m_*) divided by the total number of pairs that satisfy case 1 (2*N_i_* × 2*N_j_*). Here, the number of genes are twice as many as that of individuals because we assume diploid species. Similarly, the first term in the other five equations stand for the corresponding proportion of two-genes pairs. The second term stands for the probability of non-identical-by-descent. Considering which stages the two genes sampled belonged to in year *t* — 1, we formulate the probability by that in year *t* – 1 (i.e., *H*_*mm,t*-1_ or *H*_*kl,t*-1_) except 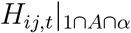. In the case of 1 ⋂ *A* ⋂ *α*, one gene is sampled from genes that moved from stage *m* to *i* by survival, while the other is from those that moved from stage *m* and *j* by survival. These two sampling sources are mutually exclusive and are not independent representatives of stage *m* of year *t* — 1, because one gene in stage *m* in year *t* — 1 could not move to both stage *i* and *j* simultaneously by survival. In other words, a gene in stage *m* in the previous year cannot be sampled twice, which violates the assumption of *H*_*mm,t*-1_, that is, “sampling with replacement.” Therefore, 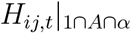 inherits the probability that two genes randomly sampled from stage m “without” replacement in year *t* – 1 were not identical-by-descent, which can be obtained by dividing *H*_*mm,t*-1_ by the chance of not sampling the same gene twice 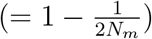.

Substituting equations 5 to equation 4, *H_ij,t_* is formulated as follows.

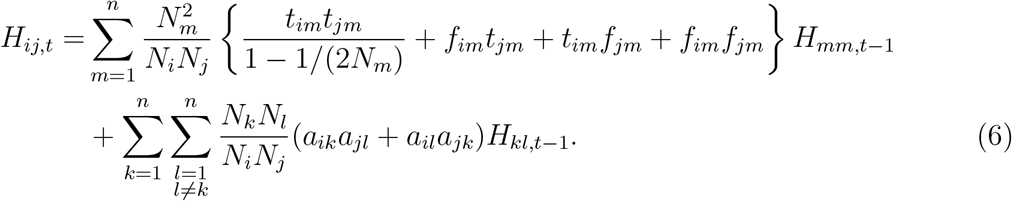

As for case 2, We decompose *H_ii,t_* into six subsets.

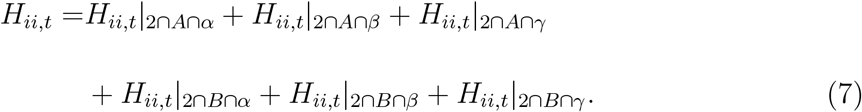

The probabilities of non-identical-by-descent on the right side of equation 7 can be formulated with *H*_*mm,t*-1_ and *H*_*kl,t*-1_, as previously done for *H_ij,t_* in case 1 (see Supporting Information 1.2).

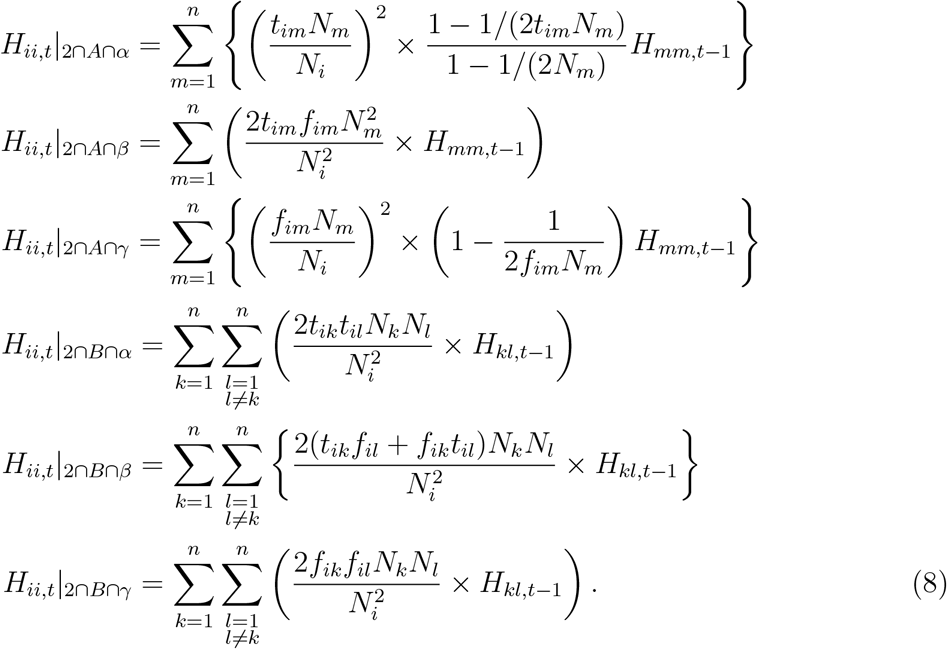

Here, the second term of 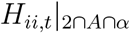 and 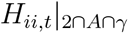 are not exactly the same as *H*_*mm,t*-1_. This is because the sources from which two genes are sampled are not independent representatives of stage *m* of the previous year and we need adjustment on *H*_*mm,t*-1_, as with the case of 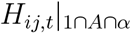. In case 2 ⋂ *A* ⋂ *α*, for example, two genes are always sampled from a common subset of stage *m*, that is, genes transferred from stage *m* to *i* by survival. Therefore, two genes are not sampled separately and independently from the whole stage m of the previous year. In case 2 ⋂ *A* ⋂ *γ*, two genes are sampled from a common source (i.e., genes that were transferred from stage m to *i* by reproduction). Reproduction allows genes to be duplicated multiple times, which deviates the gene frequencies and the chance of non-identical-by-descent of the sampling source from those of stage *m* in year *t* – 1. We adjusted *H*_*mm,t*-1_ by multiplying 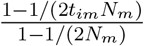 and 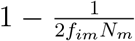. in case 2 ⋂ *A* ⋂ *α* and 2 ⋂ *A* ⋂ *γ*, respectively (see Supporting Information for details).

Substituting equations 8 to equation 7, *H_ii,t_* is formulated as follows.

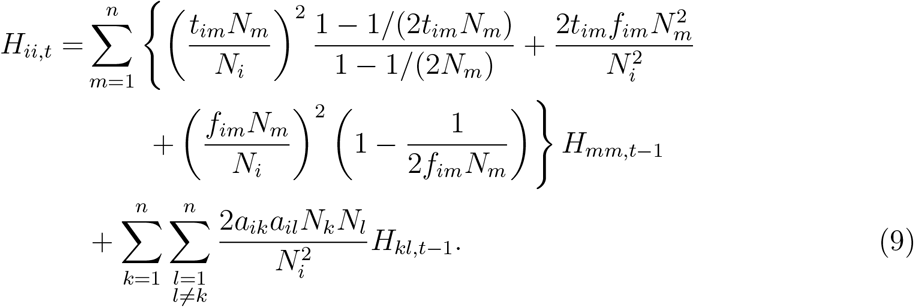

Combining case 1 (equation 6) and 2 (equation 9), we construct a matrix equation.

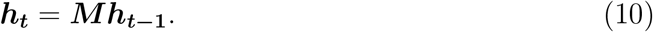

***h_t_*** and ***h_t-1_*** are vectors, each of which consists of *H_ij,t_* and *H*_*ij,t*-1_ for all possible pairs of *i* and *j* (1 ≤ *i* ≤ *n*, 1 ≤ *j* ≤ *n*). As the number of two-stage pairs is 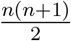, both ***h_t_*** and ***h***_***t*-1**_ have 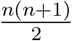 elements. ***M*** is a square matrix whose dimension is 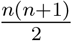 and whose elements are equal to the corresponding coefficients of *H*_*mm,t*-1_ and *H*_*kl,t*-1_ in equations 6 and 9 (see Supporting Information 2 for the detailed elements of **M**). The order of elements in ***h_t_*** is arbitrary as long as it matches with that in ***h***_***t*-1**_ and ***M***.

In general, multiplying matrix ***M*** will be asymptotically the same as multiplying the dominant eigenvalue of ***M***, while ***h_t_*** converges to a scalar multiplication of the leading right eigenvector, for sufficiently large *t*.

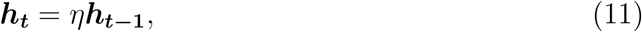

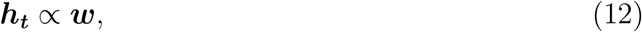

where *η* and ***w*** are the leading eigenvalue and its corresponding right eigenvector of matrix **M**, respectively. We denote *w_ij_* as the element of **w** that corresponds to *H_ij,t_* of ***h_t_***.

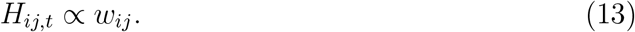

Equation 11 means that *H_ij,t_* changes with a constant rate *η* over the course of time for all *i* and *j*. Here, we denote *H_t_* as expected heterozygosity of the whole population in time *t*. *H_t_* can be formulated as the sum of *H_ij,t_* weighted by the number of individuals in stage *i* and *j*.

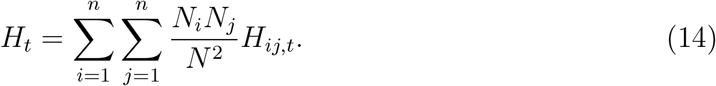

Because we assume that population size (*N*) and the number of individuals in a given stage *i* (*N_i_*) are constant, *H_t_* changes with the same rate as *H_ij,t_* that is, *η*.

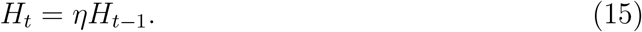

#### 2.1.3 Demographic genetic structure and effective population size

Demographic genetic structure is the comparison of neutral genetic diversity among life history stages at a particular time point. We formulate theoretical expectation of demographic genetic structure by the logarithm of the ratio of expected heterozygosity between different stages. Equation 13 indicates that the relative ratio of expected heterozygosity among stages can be approximated by comparing the elements of ***w***. With regard to the comparison between stage *i* and *j*, demographic genetic structure is formulated as follows.

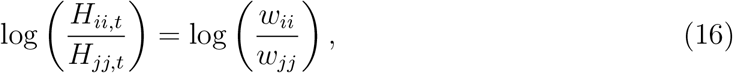

When 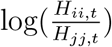 is positive, *H_ii,t_* is larger than *H_jj,t_* (genetic diversity is higher in stage *i* than in stage *j*), and when negative vice versa. It should be noted that 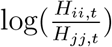 is time-invariant, although *H_ii,t_* and *H_jj,t_* themselves change with time.

We also obtain the equation for effective population size *N_e_* from our model. The probability of non-identical-by-descent of the overall population decreases with the rate of 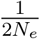 per generation time (Crow & Kimura, 1970).

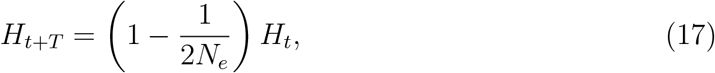

where *T* is generation time and is defined as the mean age of net fecundity in the cohort (Carey & Roach, 2020; see Supporting Information 1.3 for details). Considering that *H_t_* changes with the rate of *η* per year (equation 15), 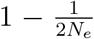 should be equivalent to *η^T^*. Therefore, We formulate *N_e_* as follows.

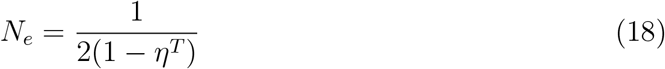

To sum up, demographic genetic structure and effective population size are derived from the leading right eigenvector and the dominant eigenvalue of matrix ***M***, respectively. Therefore, our matrix model integrates the two proxies of the temporal genetic dynamics, facilitating comprehensive understandings on demographic genetic structure.

### 2.2 Validation of the model

To ensure that the procedures of model formulation are adequate, we compared theoretically obtained *η* and demographic genetic structure with observed ones computed by stochastic simulation. Firstly, we arranged a set of life histories to be used for the comparison between theory and simulation. We considered life histories of perennial plants with two (*n* = 2: juvenile and adult) and three stages (*n* = 3: seed, juvenile, and adult; Figure 2). Equation 10 can be rewritten as follows.

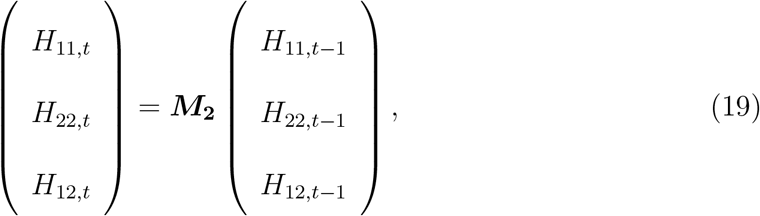

and

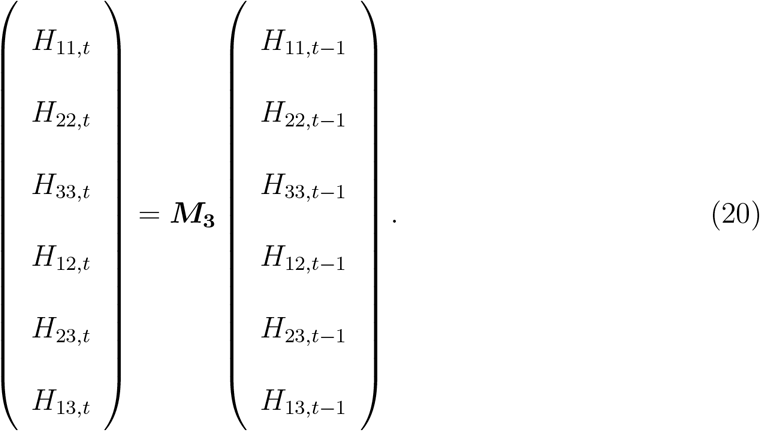

**Figure 2:**
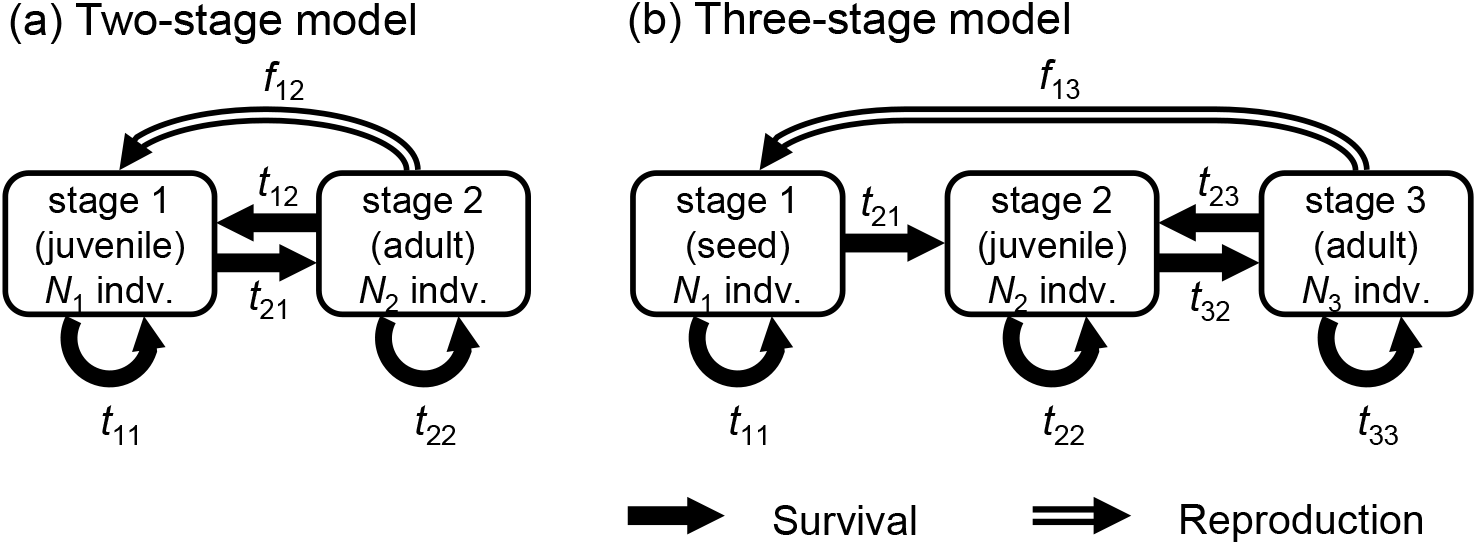
The two model used in analysis: (a) two-stage model and (b) three stage model. Arrows represent flow of individuals, or genes, either by survival (single line) or reproduction (double line)

Equation 19 and 20 correspond to the case of *n* = 2 and *n* = 3, respectively. The elements of ***M***_**2**_ and ***M***_**3**_ are functions of demographic rates (*t_ij_, f_ij_*, *a_ij_*) and the number of individuals in each stage (*N_i_*, see Supporting Information 2 for details). We determined these variables by the following procedures. Firstly, total population size *N* was set to 100, which was randomly divided into all possible flows of individuals within a population. In the case of the two-stage model, for example, 100 individuals were randomly split into five flows: stasis at juvenile, growth from juvenile to adult, retrogression from adult to juvenile, stasis at adult, and reproduction (Figure 2a). *N*_1_ and *N*_2_ were calculated as the sum of flows coming into stage 1 and 2, respectively. Demographic rates (*t*_11_, *t*_21_, *t*_12_, *t*_22_, and *f*_12_) were determined by dividing the number of individuals of corresponding flows by that of the stages from which the flows came out. Similarly, in the three-stage model, 100 individuals were divided into seven flows (Figure 2b), and the number of individuals and demographic rates were determined. Details of splitting N and determining demographic rates are explained in Figure S1. Five hundred sets of parameter values were randomly generated for both the two- and three-stage models, which covered a wide range of the parameter space (Figure S2). To consider the situation of *N* = 500 and *N* = 1,000, we multiplied *N*_1_ and *N*_2_ (when *n* = 3, *N*_3_ as well) by 5 and 10 while keeping demographic rates unchanged. In total, we considered 1,500 sets of parameter values (500 sets of demographic rates × 3 sets of *N*) for each of the two- and the three-stage model.

For each parameter set, we simulated 200 years of temporal dynamics of expected heterozygosity at a neutral biallelic locus 100 times. We calculated the mean expected heterozygosity over the 100 replicates for the overall population and all the two-stage pairs at every t, which were denoted as 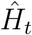 and 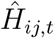, respectively. All simulations were initiated with maximum expected heterozygosity, in which two alleles share the gene pool half-and-half in all stages (i.e., *H*_0_ = *H*_*ij*,0_ = 0.5 for all *i* and *j*). We calculated the annual change rate of 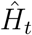 by

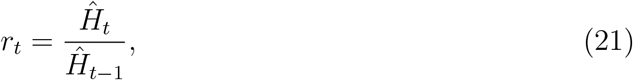

where 1 ≤ *t* ≤ 200. We took logarithm of *r_t_* and calculated its mean and standard error, which were compared to *η. η* is the theoretical counterpart *r_t_* and was obtained as the dominant eigenvalue of matrix **M_2_** or **M_3_**.

Using simulation results, we also calculated the mean of demographic genetic structure over the 200 years. As for the two-stage model, we calculated 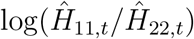. We also calculated 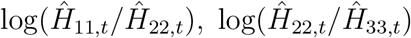 and 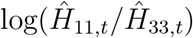 in the case of the three-stage model. These four proxies of observed demographic genetic structures were compared to theoretical counterparts, that is, log(*H*_11,*t*_/*H*_22,*t*_) for the two-stage model, as well as log(*H*_11,*t*_/*H*_22,*t*_), log(*H*_22,*t*_/*H*_33,*t*_) and log(*H*_11,*t*_/*H*_33,*t*_) for the three-stage model. These four logarithmic ratios were obtained by solving the leading right eigenvector of **M_2_** and **M_3_** and substituting their elements to equation 16.

### 2.3 Analysis on demographic genetic structure

When interpreting demographic genetic structure, previous studies compared immature (approximately young) stages with mature (approximately old) ones to predict temporal changes. To judge if such comparison could serve as a proxy for temporal dynamics of genetic diversity, we examined the four logarithmic ratios of expected heterozygosity theoretically obtained in the previous section (i.e., log(*H*_11,*t*_/*H*_22,*t*_) for the two-stage model, and log(*H*_11,*t*_/*H*_22,*t*_), log(*H*_22,*t*_/*H*_33,*t*_) and log(*H*_11,*t*_/*H*_33,*t*_) for the three-stage model). For the same 3,000 parameter sets as “Validation of the model” section, we analytically obtained *η* and *N_e_*, which reflect the change rate of expected heterozygosity per year and per generation, respectively. *η* was obtained by solving the dominant eigenvalue of ***M*_2_** and ***M*_3_**. Then, using *η*, we obtained *Ne* based on equation 18. We examined if *η* and *N_e_*, both of which genuinely represent temporal dynamics of genetic diversity, were correlated with the four logarithmic ratio that stood for demographic genetic structure.

Moreover, to explore basic behaviors of demographic genetic structure, we analyzed the dependence of demographic genetic structure on total population size *N* and stable stage distribution. Stable stage distribution was quantified by the logarithm of the ratio among *N*_1_, *N*_2_, and *N*_3_ (i.e., log(*N*_1_/*N*_2_), log(*N*_2_/*N*_3_), and log(*N*_1_/*N*_3_)).

## 3 Results

### 3.1 Validation of the model

The rate of change in expected heterozygosity of the overall populations (*r_t_*), which was computed by simulation, took almost exactly the same value as the theoretical counterpart *η* for all 500 sets of parameter values in both the two- and the three-stage models (Figure 3, S3).

**Figure 3:**
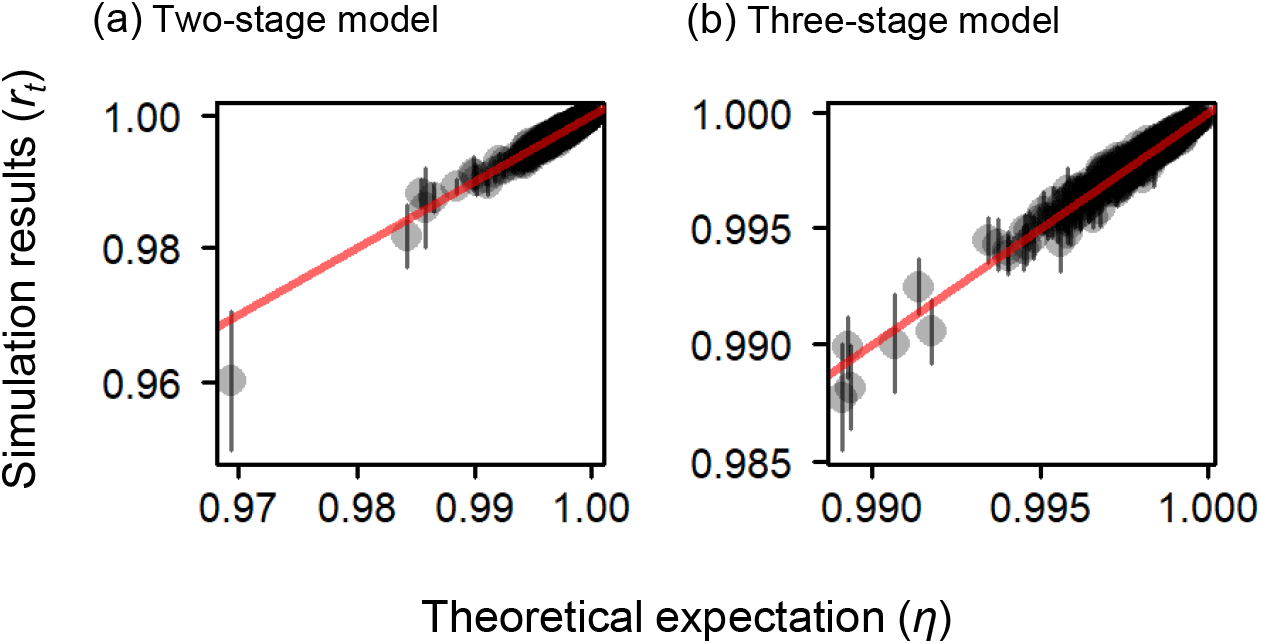
Comparison of the annual change rate of expected heterozygosity of the overall population between the theoretical expectations (*η*) and the simulation results (*r_t_*) for (a) the two-stage and (b) the three-stage model when *N* = 100. Each gray semi-transparent point corresponds to one of the 500 parameter sets. As for *r_t_*, geometric mean over 1 ≤ *t* ≤ 200 is shown with standard error (vertical bar). Red lines represent *η* = *r_t_*

Comparison of demographic genetic structure between simulation and analytical results turned out that our theoretical model yielded almost equivalent logarithmic ratio of expected heterozygosity among stages to that of simulation (Figure 4, S4, S5).

**Figure 4:**
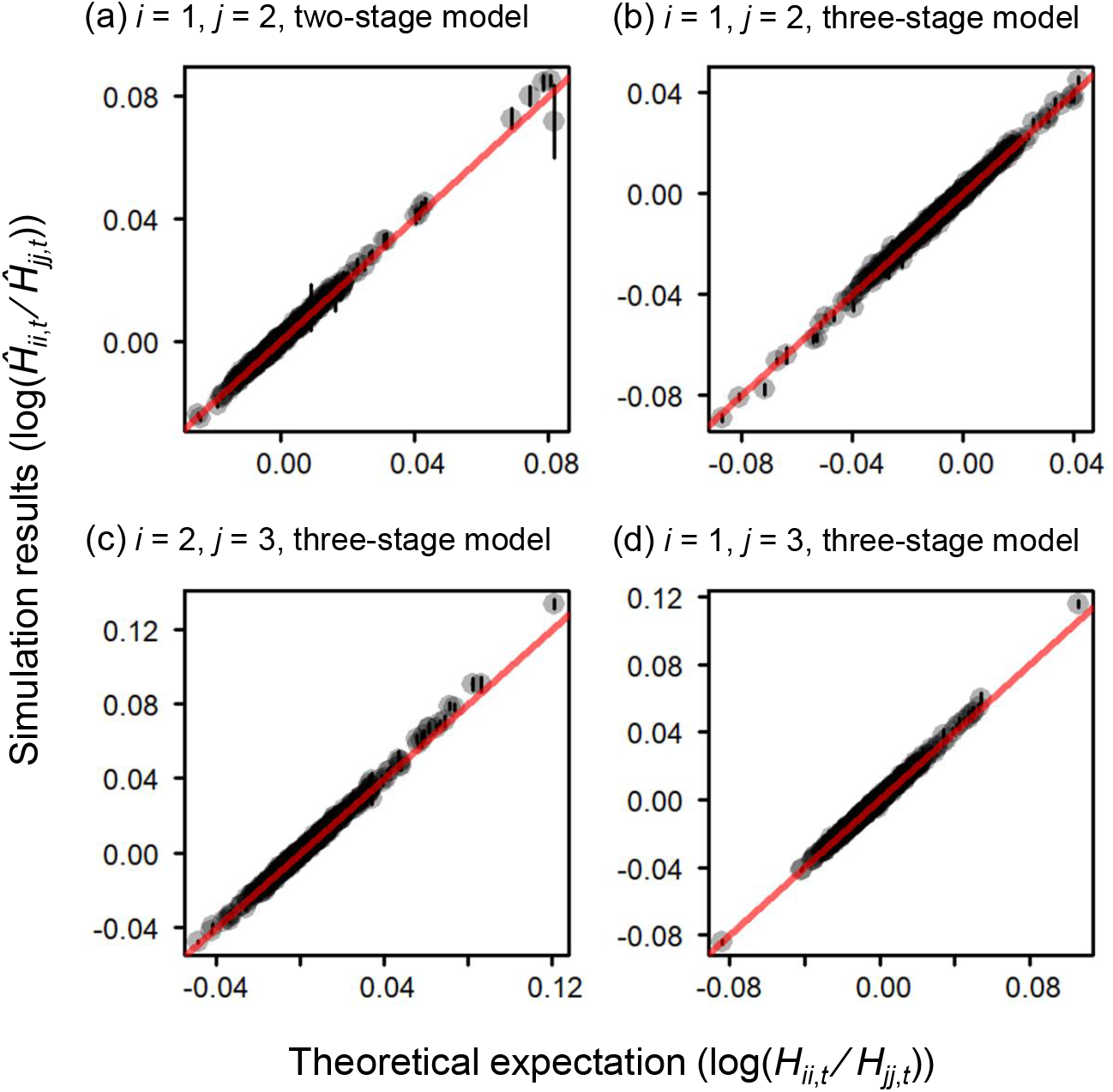
Comparison of demographic genetic structure between the theoretical expectations (log(*H_ii,t_*/*H_jj,t_*)) and the simulation results 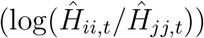 when *N* = 100. Each gray semi-transparent point corresponds to one of the 500 parameter sets. As for the simulation results, mean and standard error (vertical bar) over 1 ≤ *t* ≤ 200 are shown. There is one proxy for the two-stage model (a: *i* = 1 and *j* = 2), while there are three proxies for the three-stage model (b: *i* = 1 and *j* = 2; c: *i* = 2 and *j* = 3; d: *i* = 1 and *j* = 3). The theoretical expectations exactly match with the simulation results when plotted on the red lines

To further confirm the validity of our model, we checked the temporal dynamics of 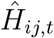 and compared it with theoretical expectation, that is, the repeated multiplication of matrix **M_2_** or **M_3_** to ***h_t_***. We found that theoretical prediction fitted well to simulation results (Figure S6).

Thus, our model seems to be valid across a wide range of parameter space.

### 3.2 Analysis on demographic genetic structure

All the four proxies of demographic genetic structure, which are theoretically obtained based on equation 16, have an apparent correlation neither with *N_e_* nor with *η* regardless of *N* (Figure 5, S7-11). On the other hand, demographic genetic structure is clearly associated with total population size *N*. As *N* increases, all the four logarithmic ratios gradually converge to zero, which means that expected heterozygosity becomes equal among stages (Figure 6). Moreover, there is a strong positive correlation with stable stage distribution: expected heterozygosity is higher in stages with more individuals (Figure 7). The correlation becomes weaker with increasing *N*, as logarithmic ratios converge to zero.

**Figure 5:**
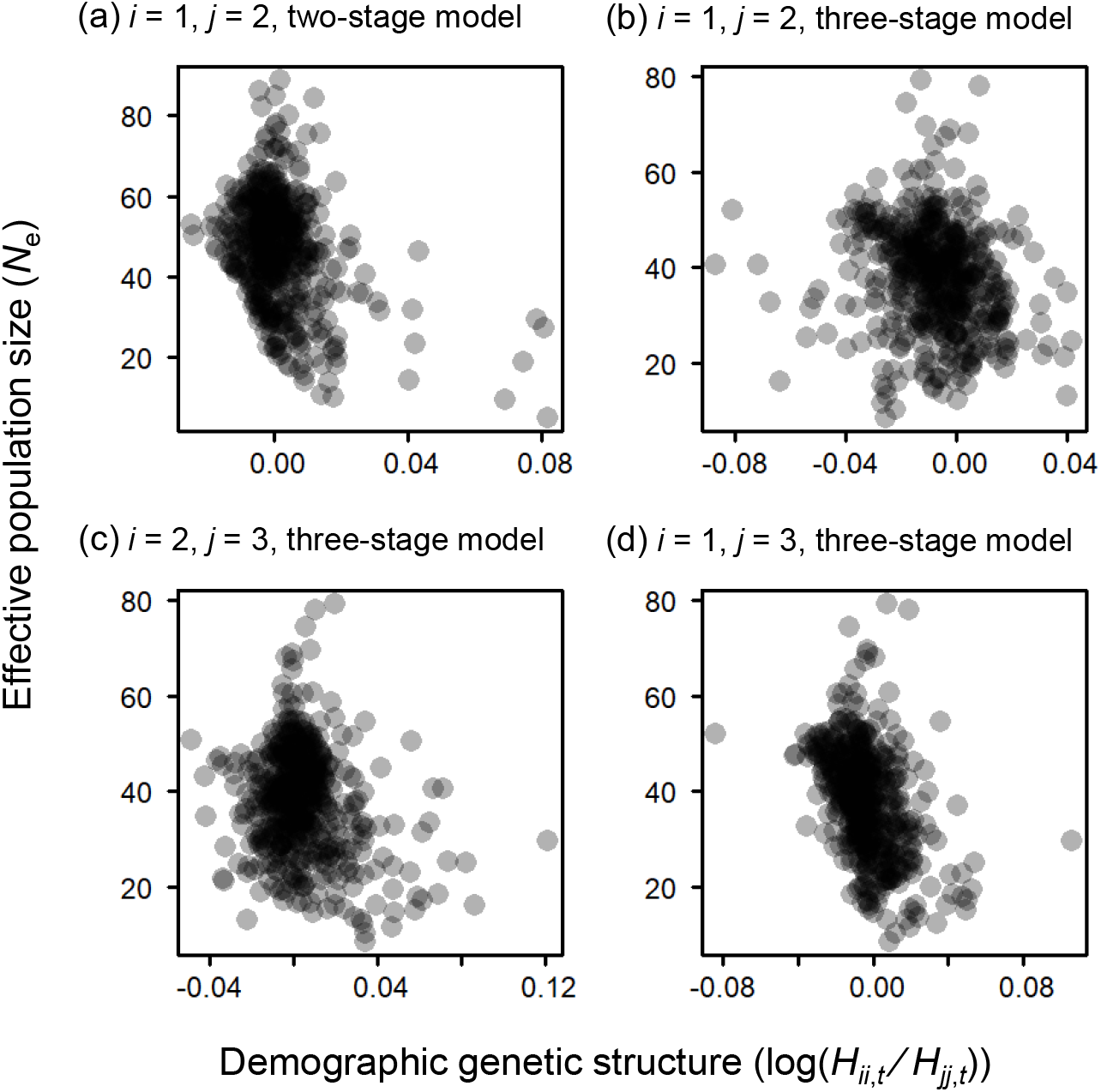
Comparison of demographic genetic structure (log(*H_ii,t_*/*H_jj,t_*)) with effective population size (*N_e_*) when *N* = 100. (a) *i* =1 and *j* = 2 of the two-stage model, (b) *i* = 1 and *j* = 2, (c) *i* = 2 and *j* = 3, (d) *i* = 1 and *j* = 3 of the three-stage model

**Figure 6:**
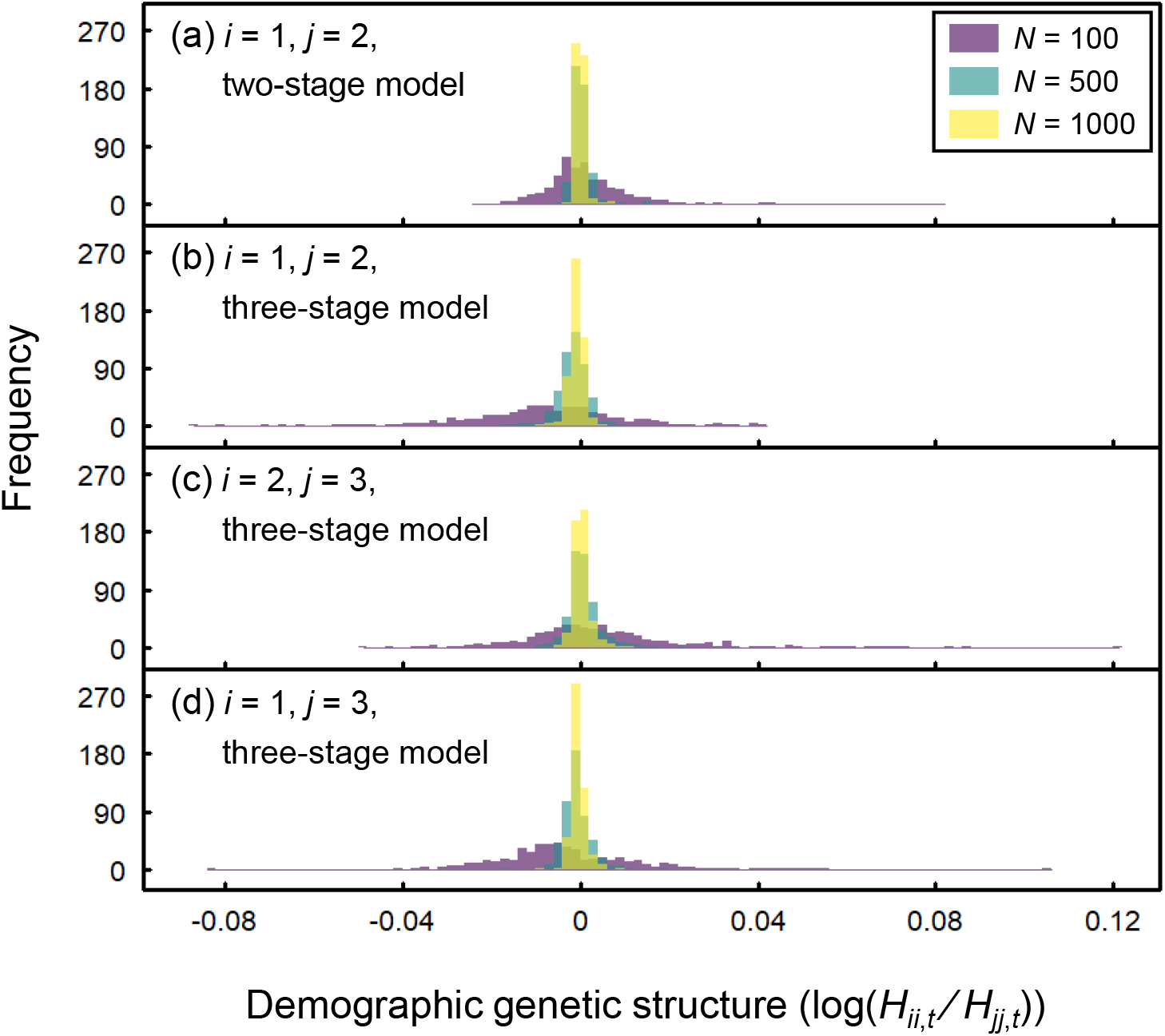
Histogram of demographic genetic structure (log(*H_ii,t_/H_jj,t_*)) with varying *N*. (a) log(*H*_11,*t*_/*H*_22,*t*_) of the two-stage model, (b) log(*H*_11,*t*_/*H*_22,*t*_), (c) log(*H*_22,*t*_/*H*_33,*t*_), (d) log(*H*_11,*t*_/*H*_33,*t*_) of the three-stage model

## 4 Discussion

### 4.1 Interpreting demographic genetic structure

A common interpretation on demographic genetic structure is that if juvenile stages are less diverse than mature stages, genetic diversity would decrease with time over the course of generation turnover. However, our model shows that relative ratio of expected heterozygosity between stage classes does not correlate with either *N_e_* or *η*: even though *N_e_* and *η* are small, expected heterozygosity does not necessarily decline from mature to juvenile stages. Therefore, inferring temporal trends in genetic diversity solely from demographic genetic structure is potentially misleading. This study, to our knowledge, for the first time draws caution on the conventional use of demographic genetic structure.

Many previous empirical studies that analyzed demographic genetic structure found that genetic diversity did not decrease from the most mature to the most immature stages and took comparable values among stages (Aldrich et al., 1998; Ally & Ritland, 2007; Linhart et al., 1981; Murren, 2003; Schmidt et al., 2018; Vranckx et al., 2014). Our model shows that the logarithmic ratio of expected heterozygosity is distributed around zero, especially under large N, indicating that expected heterozygosity is basically almost equivalent to one another. Therefore, our model might be in line with previous empirical results.

While demographic genetic structure is irrelevant to temporal dynamics, it is tightly linked to stable stage distribution: expected heterozygosity is relatively high in stage with more individuals, and low in stage with less individuals (Figure 6). In general, small number of individuals intensifies stochastic genetic drift due to increased sampling bias in gene frequencies, leading to the loss of genetic diversity (Kimura and Crow 1970). When stage distribution is skewed, the degree of stochasticity will vary among stages. Stage with smaller number of individuals is made up of genes that were sampled fewer times from the gene pool of the previous year, thus suffering random perturbation in gene frequencies to a greater extent. The alleviated stochasticity must have resulted in the lower genetic diversity in stages with fewer individuals.

As the total population size *N* increased, inter-stage difference in genetic diversity disappears even under the skewed stage distribution (Figure 7). This result indicates that the number of individuals of each stage is large enough to reduce stochasticity under large *N*, leading to comparable level of genetic diversity among stages. Large population size also contributes to the maintenance of genetic diversity, because *N_e_* increases and η approaches to 1 with increasing *N* (Figure S12).

**Figure 7:**
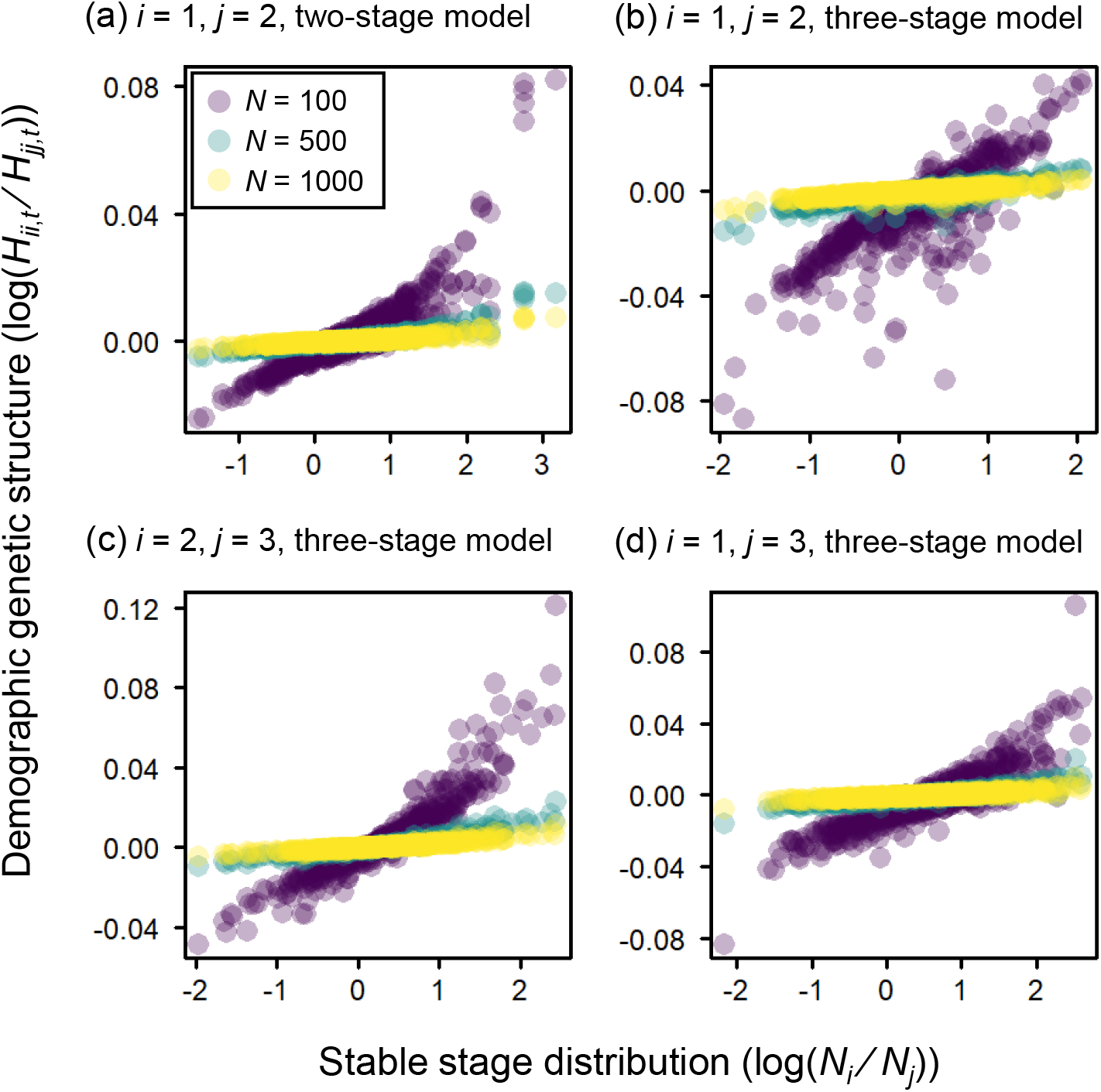
Relationships between stable stage distribution (log(*N_i_*/*N_j_*)) and demographic genetic structure (log(*H_ii,t_*/*H_jj,t_*)) with varying *N*.

To sum up, it can be said that genetic diversity becomes uneven among life history stages under small population size and that the unevenness among stages reflects stable stage distribution rather than the temporal dynamics of genetic diversity.

### 4.2 Future application of our model

Our model not only provides theoretical background of demographic genetic structure, but also has some potential for application. One possible application is to compare raw demographic genetic structure, which is obtained by any neutral genetic markers, with the theoretical expectation calculated based on the equations we derived. The deviations of observed structure from expectation reflect factors unexplored in our model, such as fluctuating population size, non-random mating, selection, and immigration. Thus, our model can work as a null model of demographic genetic structure. To make the most use of our equations, it is necessary to monitor individuals from year to year to estimate demographic rates of each stage class. If long-term demographic monitoring is unavailable or impractical for some reasons, recording relative number of individuals among stage classes at a single time point would be at least desirable to consider stage distribution, which turned out to be a major determinant of demographic genetic structure in our model.

In addition to demographic genetic structure, we develop a new statistics *η*, which is the annual change rate of expected heterozygosity, which can be potentially useful for population viability assessment. Whether population size can be maintained over time (i.e., population growth rate remains high) is considered as a criterion of long-term population persistence (Hens et al., 2017; Knight et al., 2009). Demographic rates have been used to calculate population growth rate per year (usually denoted as *λ*) by solving the eigenvalue problem of matrix population models (equation 3) (Caswell, 2001; Crone et al., 2011). While it is acknowledged that not only population size but also genetic diversity should be maintained for long-term population viability, there has been no counterpart of population growth rate that can evaluate the change rate of genetic diversity per year (not per generation time). Being a change rate per year, *η* is directly linked to temporal change in genetic diversity compared to demographic genetic structure and effective population size, and enables us to assess genetic diversity at the same time scale as population dynamics. Therefore, *η* can serve as the counterpart of *λ* and can be an useful proxy to evaluate population viability from genetic point of view. Although we assume perennial plants in this study, our model can be applied to other taxa that are structured by stage. Moreover, because age structure, in which the probabilities of stasis and retrogression are zero, is a special case of stage structure, our model can also be applied to populations with age structure. Therefore, evaluating η for a variety types of structured populations will be a future step to make the best use of our model.

## Acknowledgements

This research was financially supported by Grands-in-Aid for Scientific Research from the JSPS KAKENHI (Grant no. 19H03294, 20K06821, and 21J10814).

## Author Contribution

Y.T. conceived the idea, Y.T. and T.T. developed and analyze the model, Y.T., T.T., and M.O. wrote the manuscript. All the authors contributed critically to the drafts and gave final approval for publication.

## Data Availability Statement

We intend archive our data and R scripts at Dryad Digital Repository when this manuscript is accepted.

## References

Agashe, D., Falk, J. J., & Bolnick, D. I. (2011). Effects of founding genetic variation on adaptation to a novel resource. Evolution, 65(9), 2481–2491. https://doi.org/10.1111/j.1558-5646.2011.01307.x

Aldrich, P. R., Hamrick, J. L., Chavarriaga, P., & Kochert, G. (1998). Microsatellite analysis of demographic genetic structure in fragmented populations of the tropical tree *Symphonia globulifera*. Molecular Ecology, 7, 933–944. https://doi.org/10.1046/j.1365-294x.1998.00396.x

Ally, D., & Ritland, K. (2007). A case study: looking at the effects of fragmentation on genetic structure in different life history stages of old-growth mountain hemlock *(Tsuga mertensiana)*. Journal of Heredity, 98(1), 73–78. https://doi.org/10.1093/jhered/esl048

Barrett, R. D. H., & Schluter, D. (2008). Adaptation from standing genetic variation. Trends in Ecology and Evolution, 23(1), 38–44. https://doi.org/10.1016/j.tree.2007.09.008

Carey, J. R., & Roach, D. A. (2020). Biodemography: An Introduction to Concepts and Methods. Princeton University Press.

Caswell, H. (2001). Matrix population models (2nd edn.). Sinauer Associates, Inc.

Charlesworth, B. (2008). Evolution in age-structured populations (2nd edn.). Cambridge University Press.

Crone, E. E., Menges, E. S., Ellis, M. M., Bell, T., Bierzychudek, P., Ehrlén, J., Kaye, T. N., Knight, T. M., Lesica, P., Morris, W. F., Oostermeijer, G., Quintana-Ascencio, P. F., Stanley, A., Ticktin, T., Valverde, T., & Williams, L. L. (2011). How do plant ecologists use matrix population models? Ecology Letters, 14(1), 1–8. https://doi.org/10.1111/j.1461-0248.2010.01540.x

Crow, J. F., & Kimura, M. (1970). An introduction in population genetic theory. Harper and Row.

Felsenstein, J. (1971). Inbreeding and variance effective numbers in populations with overlapping generations. Genetics, 68, 581–597. https://doi.org/10.1093/genetics/68.4.581

Hens, H., Pakanen, V., Jäkäläniemi, A., Tuomi J., & Kvist, L. (2017). Low population viability in small endangered orchid populations: Genetic variation, seedling recruitment and stochasticity. Biological Conservation, 210, 174–183. https://doi.org/10.1016/j.biocon.2017.04.019

Hill, W. G. (1972). Effective size of populations with overlapping generations. Theoretical Population Biology, 3, 278–289. https://doi.org/10.1016/0040-5809(72)90004-4

Hill, W. G. (1979). A note on effective population size with overlapping generations. Genetics, 92(1), 317–322. https://doi.org/10.1093/genetics/92.1.317

Johnson, D. L. (1977). Inbreeding in populations with overlapping generations. Genetics, 87(3), 581–591. https://doi.org/10.1093/genetics/87.3.581

Kettle, D. J., Hollingsworth, P. M., Jaffré, T., Moran, B., & Ennos, R. A. (2007). Identifying the early genetic consequences of habitat degradation in a highly threatened tropical conifer, *Araucaria nemorosa* Laubenfels. Molecular Ecology, 16, 3581–3591. https://doi.org/10.1111/j.1365-294X.2007.03419.x

Knight, T. M. (2004). The effects of herbivory and pollen limitation on a declining population of *Trillium grandiflorum*. Ecological Applications, 14(3), 915–928. https://doi.org/10.1890/03-5048

Knight, T. M., Caswell, H., & Kalisz, S. (2009). Population growth rate of a common understory herb decreases non-linearly across a gradient of deer herbivory Forest Ecology and Management, 257(3), 1095–1103. https://doi.org/10.1016/j.foreco.2008.11.018

Linhart, Y. B., Mitton, J. B., Sturgeon, K. B., & Davis, M. L. (1981). Genetic variation in space and time in a population of ponderosa pine. Heredity, 46(3), 407–426. https://doi.org/10.1038/hdy.1981.49

Mimura, M., Yahara, T., Faith, D. P., Vázquez-Domínguez, E., Colautti, R. I., Araki, H., Javadi, F., Núñez-Farfán, J., Mori, A. S., Zhou, S., Hollingsworth, P. M., Neaves, L. E., Fukano, Y., Smith, G. F., Sato, Y., Tachida, H., & Hendry, A. P. (2017). Understanding and monitoring the consequences of human impacts on intraspecific variation. Evolutionary Application, 10, 121–139. https://doi.org/10.1111/eva.12436

Murren, C. J. (2003). Spatial and demographic population genetic structure in *Catase-tum viridiflavum* across a human-disturbed habitat. Journal of Evolutionary Biology, 16(2), 333–342. https://doi.org/10.1046/j.1420-9101.2003.00517.x

Ohara, M., Takada, T., & Kawano, S. (2001). Demography and reproductive strategies of a polycarpic perennial, *Trillium apetalon* (Trilliaceae). Plant Species Biology, 16(3), 209–217. https://doi.org/10.1046/j.1442-1984.2001.00062.x

Orive, M. E. (1993). Effective population size in organisms with complex life-histories. Theoretcal Population Biology, 44, 316–340. https://doi.org/10.1006/tpbi.1993.1031

Ramsayer, J., Kaltz, O. & Hochberg, M. E. (2012). Evolutionary rescue in populations of *Pseudomonas flurescens* across an antibiotic gradient. Evolutionary Application, 6, 608–616. https://doi.org/10.1111/eva.12046

Schmidt, D. J., Fallon, S., Roberts, D. T., Espinoza, T., McDougall, A., Brooks, S. G., Kind, P. K., Bond, N. R., Kennard, M. J., Hughes, J. M. (2018). Monitoring age-related trends in genomic diversity of Australian lungfish. Molecular Ecology, 27(16), 3231–3241. https://doi.org/10.1111/mec.14791

Silvertown, J. (1987). Introduction to plant population ecology (2nd edn). Longman Scientific & Technical.

Tomimatsu, H., & Ohara, M. (2010). Demographic response of plant populations to habitat fragmentation and temporal environmental variability. Oecologia, 162, 903–911. https://doi.org/10.1007/s00442-009-1505-8

Torices, R., Méndez, M., & Gómez, J. M. (2011). Where do monomorphic sexual systems fit in the evolution of dioecy? Insights from the largest family of angiosperms. New Phytologist, 190(1), 234–248. https://doi.org/10.1111/j.1469-8137.2010.03609.x

Tsuzuki, Y., Sato, M. P., Matsuo, A., Suyama, Y., & Ohara, M. (2021). Genetic consequences of habitat fragmentation in a perennial plant *Trillium camschatcense* are subjected to its slow-paced life history. Population Ecology, 64(1), 5–18. https://doi.org/10.1002/1438-390X.12093

Vranckx, G., Jacquemyn, H., Mergeay, J., Cox, K. Kint, V., & Honnay, O. (2014). Transmission of genetic variation from the adult generation to naturally established seedling cohorts in small forest stands of pedunculate oak *(Quercus robur* L.). Forest Ecology and Management, 312, 19–27. https://doi.org/10.1016/j.foreco.2013.10.027

Waples, R. S., Do, C., & Chopelet, J. (2011). Calculating Ne and Ne/N in age-structured populations: a hybrid Felsenstein-Hill approach. Ecology, 92(7), 1513–1522. https://doi.org/10.1890/10-1796.1

Waples, R. S., Luikart, G., Faulkner, J. R., & Tallmon, D. A. (2013). Simple life-history traits explain key effective population size ratios across diverse taxa. Proceedings of the Royal Society B, 280(1768), 20131339. https://doi.org/10.1098/rspb.2013.1339

Yonezawa, K., Kinoshita, E., Watano, Y., & Zentoh, H. (2000). Formulation and estimation of the effective population size of stage-structured populations in *Fritillaria camtschatcensis*, a perennial herb with a complex life history. Evolution, 54(6), 2007–2013. https://doi.org/10.1111/j.0014-3820.2000.tb01244.x

