## Supporting Information for "Modelling temporal dynamics of genetic diversity in stage-structured plant populations with reference to demographic genetic structure"

N10W5, Kita-ku, Sapporo City, Hokkaido Prefecture, Japan, 060-0810

### 1 Model development

#### 1.1 Formulation of $H_{ij,t}$

As explained in the main text, $H_{ij,t}$ is split into six subsets:

$$H_{ij,t} = H_{ij,t}|_{1\cap A\cap\alpha} + H_{ij,t}|_{1\cap A\cap\beta} + H_{ij,t}|_{1\cap A\cap\gamma} + H_{ij,t}|_{1\cap B\cap\alpha} + H_{ij,t}|_{1\cap B\cap\beta} + H_{ij,t}|_{1\cap B\cap\gamma}, \tag{S1}$$

where $H_{ij,t}|_{X\cap Y\cap Z}$ stands for $H_{ij,t}$ under the concurrence of case X, Y, and Z ($X = 1, 2$; $Y = A, B$; $Z = \alpha, \beta, \gamma$).

We define sub-stage $i_{ms}$ and $i_{mr}$, which consist of individuals transferred from stage $m$ to $i$ by survival and by reproduction, respectively. Each $H_{ij,t}|_{1\cap A\cap Z}$ can be formulated using $H_{i_{ms}j_{ms},t}$ (case $1 \cap A \cap \alpha$), $H_{i_{ms}j_{mr},t}$, $H_{i_{mr}j_{ms},t}$ (case $1 \cap A \cap \beta$), and $H_{i_{mr}j_{mr},t}$ (case $1 \cap A \cap \gamma$) weighted by the corresponding number of two-gene pairs.

$$H_{ij,t}|_{1\cap A\cap\alpha} = \sum_{m=1}^{n} \frac{N_{i_{ms}}N_{j_{ms}}}{N_iN_j} H_{i_{ms}j_{ms},t} \tag{S2}$$

$$H_{ij,t}|_{1\cap A\cap\beta} = \sum_{m=1}^{n} \left( \frac{N_{i_{ms}}N_{j_{mr}}}{N_iN_j} H_{i_{ms}j_{mr},t} + \frac{N_{i_{mr}}N_{j_{ms}}}{N_iN_j} H_{i_{mr}j_{ms},t} \right) \tag{S3}$$

$$H_{ij,t}|_{1\cap A\cap\gamma} = \sum_{m=1}^{n} \frac{N_{i_{mr}}N_{j_{mr}}}{N_iN_j} H_{i_{mr}j_{mr},t}. \tag{S4}$$

Here, $N_{i_{ms}}$ and $N_{j_{ms}}$ denote the number of individuals in sub-stage $i_{ms}$ and $j_{ms}$, respectively. As for case $1 \cap A \cap \alpha$, two genes, each sampled from stage $i$ and $j$, belong to sub-stage $i_{ms}$ and $j_{ms}$ with the chance of $\frac{2N_{i_{ms}}}{2N_i} \times \frac{2N_{j_{ms}}}{2N_j}$. The number of genes is twice as many as that of individuals because we assume diploid species. Thus, $H_{i_{ms}j_{ms},t}$ is weighted by $\frac{N_{i_{ms}}N_{j_{ms}}}{N_iN_j}$, as shown in equation S2 . Case $1 \cap A \cap \beta$ (equation S3 ) and $1 \cap A \cap \gamma$ ( equation S4 ) are similarly formulated.

In the concurrence of case 1, A, and $\alpha$, two genes, each sampled from sub-stage $i_{ms}$ and $j_{ms}$, cannot be the same gene because one gene in stage $m$ in year $t-1$ could not move to both stage $i$ and $j$ simultaneously by survival. Therefore, $H_{i_{ms}j_{ms},t}$ is equal to the probability that two genes randomly sampled from stage $m$ “without” replacement in time $t-1$ are not identical-by-descent.

Here, we define $H'_{ij,t}$ as the probability that two genes sampled from stage $i$ and $j$ "without" replacement in time $t$ are non-identical-by-descent. Because $H_{i_{ms}j_{ms},t}$ is equal to $H'_{mm,t-1}$ in case $1 \cap A \cap \alpha$, we formulate $H'_{mm,t-1}$. When sampling two genes from stage $m$ with replacement in year $t-1$, the same gene can be sampled twice with the probability of $\frac{1}{2N_m} \times \frac{1}{2N_m} \times 2N_m = \frac{1}{2N_m}$, which makes no contribution to $H_{mm,t-1}$. Therefore, $H_{mm,t-1}$ can be formulated as follows.

$$H_{mm,t-1} = \frac{1}{2N_m} \times 0 + \left(1 - \frac{1}{2N_m}\right) \times H'_{mm,t-1} \tag{S5}$$

As a result, $H'_{mm,t-1}$ is obtained.

$$H_{i_{ms}j_{ms},t} = H'_{mm,t-1} = \frac{1}{1 - 1/(2N_m)} H_{mm,t-1} \tag{S6}$$

Unlike transfer by survival, transfer by reproduction allows the same gene to move multiple pathways simultaneously, because genes are replicated. In the case of $1 \cap A \cap \beta$ and $1 \cap A \cap \gamma$, sampling in year $t$ does not preclude the chance of sampling the same gene twice, because at least one of the two genes are transferred by reproduction. Therefore,

$$H_{i_{ms}j_{mr},t} = H_{i_{mr}j_{ms},t} = H_{i_{mr}j_{mr},t} = H_{mm,t-1} \tag{S7}$$

The number of genes in each sub-stage is given by

$$N_{i_{ms}} = t_{im} N_m \tag{S8}$$

$$N_{j_{ms}} = t_{jm} N_m \tag{S9}$$

$$N_{i_{mr}} = f_{im} N_m \tag{S10}$$

$$N_{j_{mr}} = f_{jm} N_m. \tag{S11}$$

Substituting equations S6 -S11 to equations S2 -S4 ,

$$H_{ij,t}|_{1 \cap A \cap \alpha} = \sum_{m=1}^{n} \left\{ \frac{t_{im} t_{jm} N_m^2}{N_i N_j} \times \frac{1}{1 - 1/(2N_m)} H_{mm,t-1} \right\} \tag{S12}$$

$$H_{ij,t}|_{1 \cap A \cap \beta} = \sum_{m=1}^{n} \left\{ \frac{(t_{im} f_{jm} + f_{im} t_{jm}) N_m^2}{N_i N_j} \times H_{mm,t-1} \right\} \tag{S13}$$

$$H_{ij,t}|_{1 \cap A \cap \gamma} = \sum_{m=1}^{n} \left( \frac{f_{im} f_{jm} N_m^2}{N_i N_j} \times H_{mm,t-1} \right). \tag{S14}$$

As for $H_{ij,t}|_{1 \cap B \cap Z}$ $(Z = \alpha, \beta, \gamma)$, two genes, each sampled from stage $i$ and $j$ in time $t$, were in stage $k$ and $l$, or in stage $l$ and $k$, in year $t-1$ respectively. In either situation, the probability of non-identical-by-descent remains the same as that in year $t-1$, which is $H_{kl,t-1}$, regardless of whether they were transferred only by survival (case $1 \cap B \cap \alpha$), both by survival and by reproduction (case $1 \cap B \cap \beta$), or only by reproduction (case $1 \cap B \cap \gamma$).

$$
\begin{aligned}
H_{ij,t}|_{1\cap B\cap\alpha} &= \sum_{k=1}^{n}\sum_{\substack{l=1\\ l\neq k}}^{n}\left(\frac{N_{i_{ks}}N_{j_{ls}}}{N_iN_j}H_{i_{ks}j_{ls},t}+\frac{N_{i_{ls}}N_{j_{ks}}}{N_iN_j}H_{i_{ls}j_{ks},t}\right)\\
&= \sum_{k=1}^{n}\sum_{\substack{l=1\\ l\neq k}}^{n}\left(\frac{t_{ik}t_{jl}N_kN_j}{N_iN_j}H_{kl,t-1}+\frac{t_{il}t_{jk}N_kN_j}{N_iN_j}H_{kl,t-1}\right)\\
&= \sum_{k=1}^{n}\sum_{\substack{l=1\\ l\neq k}}^{n}\left\{\frac{(t_{ik}t_{jl}+t_{il}t_{jk})N_kN_j}{N_iN_j}H_{kl,t-1}\right\} \qquad \text{(S15)}\\
H_{ij,t}|_{1\cap B\cap\beta} &= \sum_{k=1}^{n}\sum_{\substack{l=1\\ l\neq k}}^{n}\left(\frac{N_{i_{ks}}N_{j_{lr}}}{N_iN_j}H_{i_{ks}j_{lr},t}+\frac{N_{i_{kr}}N_{j_{ls}}}{N_iN_j}H_{i_{kr}j_{ls},t}\right.\\
&\qquad \left.+\frac{N_{i_{ls}}N_{j_{kr}}}{N_iN_j}H_{i_{ls}j_{kr},t}+\frac{N_{i_{lr}}N_{j_{ks}}}{N_iN_j}H_{i_{lr}j_{ks},t}\right)\\
&= \sum_{k=1}^{n}\sum_{\substack{l=1\\ l\neq k}}^{n}\left(\frac{t_{ik}f_{jl}N_kN_l}{N_iN_j}H_{kl,t-1}+\frac{f_{ik}t_{jl}N_kN_l}{N_iN_j}H_{kl,t-1}\right.\\
&\qquad \left.+\frac{t_{il}f_{jk}N_kN_l}{N_iN_j}H_{kl,t-1}+\frac{f_{il}t_{jk}N_kN_l}{N_iN_j}H_{kl,t-1}\right)\\
&= \sum_{k=1}^{n}\sum_{\substack{l=1\\ l\neq k}}^{n}\left\{\frac{(t_{ik}f_{jl}+f_{ik}t_{jl}+t_{il}f_{jk}+f_{il}t_{jk})N_kN_l}{N_iN_j}H_{kl,t-1}\right\} \qquad \text{(S16)}\\
H_{ij,t}|_{1\cap B\cap\gamma} &= \sum_{k=1}^{n}\sum_{\substack{l=1\\ l\neq k}}^{n}\left(\frac{N_{i_{kr}}N_{j_{lr}}}{N_iN_j}H_{i_{kr}j_{lr},t}+\frac{N_{i_{lr}}N_{j_{kr}}}{N_iN_j}H_{i_{lr}j_{kr},t}\right)\\
&= \sum_{k=1}^{n}\sum_{\substack{l=1\\ l\neq k}}^{n}\left(\frac{f_{ik}f_{jl}N_kN_j}{N_iN_j}H_{kl,t-1}+\frac{f_{il}f_{jk}N_kN_j}{N_iN_j}H_{kl,t-1}\right)\\
&= \sum_{k=1}^{n}\sum_{\substack{l=1\\ l\neq k}}^{n}\left\{\frac{(f_{ik}f_{jl}+f_{il}f_{jk})N_kN_j}{N_iN_j}H_{kl,t-1}\right\} \qquad \text{(S17)}
\end{aligned}
$$

Substituting equations S12 -S17 to equation S1 , we can formulate $H_{ij,t}$ as follows.

$$
\begin{aligned}
H_{ij,t} = \sum_{m=1}^{n}\left\{\frac{t_{im}t_{jm}N_m^2}{N_iN_j}\times\frac{1}{1-1/(2N_m)}+\frac{(t_{im}f_{jm}+f_{im}t_{jm})N_m^2}{N_iN_j}\right.\\
\left.\frac{f_{im}f_{jm}N_m^2}{N_iN_j}\right\}H_{mm,t-1}
\end{aligned}
$$

$$+\sum_{k=1}^{n}\sum_{\substack{l=1\\l\neq k}}^{n}\left\{\frac{(t_{ik}t_{jl}+t_{il}t_{jk})N_kN_l}{N_iN_j}+\frac{(t_{ik}f_{jl}+f_{ik}t_{jl}+t_{il}f_{jk}+f_{il}t_{jk})N_kN_l}{N_iN_j}\right.$$

$$\left.+\frac{(f_{ik}f_{jl}+f_{il}f_{jk})N_kN_l}{N_iN_j}\right\}H_{kl,t-1}$$

$$=\sum_{m=1}^{n}\frac{N_m^2}{N_iN_j}\left\{\frac{t_{im}t_{jm}}{1-1/(2N_m)}+f_{im}t_{jm}+t_{im}f_{jm}+f_{im}f_{jm}\right\}H_{mm,t-1}$$

$$+\sum_{k=1}^{n}\sum_{\substack{l=1\\l\neq k}}^{n}\frac{N_kN_l}{N_iN_j}\left((t_{ik}+f_{ik})(t_{jl}+f_{jl})+(t_{il}+f_{il})(t_{jk}+f_{jk})\right)H_{kl,t-1}.$$

$$=\sum_{m=1}^{n}\frac{N_m^2}{N_iN_j}\left\{\frac{t_{im}t_{jm}}{1-1/(2N_m)}+f_{im}t_{jm}+t_{im}f_{jm}+f_{im}f_{jm}\right\}H_{mm,t-1}$$

$$+\sum_{k=1}^{n}\sum_{\substack{l=1\\l\neq k}}^{n}\frac{N_kN_l}{N_iN_j}(a_{ik}a_{jl}+a_{il}a_{jk})H_{kl,t-1}. \qquad \text{(S18)}$$

#### 1.2 Formulation of $H_{ii,t}$

$H_{ii,t}$ is split into mutually exclusive six subsets:

$$H_{ii,t}=H_{ii,t}|_{2\cap A\cap\alpha}+H_{ii,t}|_{2\cap A\cap\beta}+H_{ii,t}|_{2\cap A\cap\gamma}$$

$$+H_{ii,t}|_{2\cap B\cap\alpha}+H_{ii,t}|_{2\cap B\cap\beta}+H_{ii,t}|_{2\cap B\cap\gamma}, \qquad \text{(S19)}$$

Considering which sub-stages two genes are sampled from, we can formulate the six $H_{ii,t}$ on the right side of equation S19 .

$$H_{ii,t}|_{2\cap A\cap\alpha}=\sum_{m=1}^{n}\left\{\left(\frac{N_{i_{ms}}}{N_i}\right)^2H_{i_{ms}i_{ms},t}\right\}$$

$$=\sum_{m=1}^{n}\left\{\left(\frac{t_{im}N_m}{N_i}\right)^2H_{i_{ms}i_{ms},t}\right\} \qquad \text{(S20)}$$

$$H_{ii,t}|_{2\cap A\cap\beta}=\sum_{m=1}^{n}\left(\frac{N_{i_{ms}}N_{i_{mr}}}{N_i^2}H_{i_{ms}i_{mr},t}+\frac{N_{i_{mr}}N_{i_{ms}}}{N_i^2}H_{i_{mr}i_{ms},t}\right)$$

$$=\sum_{m=1}^{n}\left(\frac{2N_{i_{ms}}N_{i_{mr}}}{N_i^2}H_{i_{ms}i_{mr},t}\right)$$

$$=\sum_{m=1}^{n}\left(\frac{2t_{im}f_{im}N_m^2}{N_i^2}H_{i_{ms}i_{mr},t}\right) \qquad \text{(S21)}$$

$$H_{ii,t}|_{2\cap A\cap\gamma}=\sum_{m=1}^{n}\left\{\left(\frac{N_{i_{mr}}}{N_i}\right)^2H_{i_{mr}i_{mr},t}\right\}$$

$$= \sum_{m=1}^{n} \left\{ \left( \frac{f_{im}N_m}{N_i} \right)^2 H_{i_{mr}i_{mr},t} \right\} \tag{S22}$$

$$H_{ii,t}|_{2\cap B\cap\alpha} = \sum_{k=1}^{n} \sum_{\substack{l=1 \\ l\neq k}}^{n} \left( \frac{N_{i_{ks}}N_{i_{ls}}}{N_i^2} H_{i_{ks}j_{ls},t} + \frac{N_{i_{ls}}N_{i_{ks}}}{N_i^2} H_{i_{ls}j_{ks},t} \right)$$

$$= \sum_{k=1}^{n} \sum_{\substack{l=1 \\ l\neq k}}^{n} \left( \frac{2N_{i_{ls}}N_{i_{ks}}}{N_i^2} H_{i_{ls}j_{ks},t} \right)$$

$$= \sum_{k=1}^{n} \sum_{\substack{l=1 \\ l\neq k}}^{n} \left( \frac{2t_{ik}t_{il}N_kN_l}{N_i^2} H_{i_{ls}j_{ks},t} \right) \tag{S23}$$

$$H_{ij,t}|_{2\cap B\cap\beta} = \sum_{k=1}^{n} \sum_{\substack{l=1 \\ l\neq k}}^{n} \left( \frac{N_{i_{ls}}N_{i_{kr}}}{N_i^2} H_{i_{kr}i_{ls},t} + \frac{N_{i_{kr}}N_{i_{ls}}}{N_i^2} H_{i_{kr}i_{ls},t} \right.$$

$$\left. + \frac{N_{i_{ks}}N_{i_{lr}}}{N_i^2} H_{i_{ks}i_{lr},t} + \frac{N_{i_{lr}}N_{i_{ks}}}{N_i^2} H_{i_{ks}i_{lr},t} \right)$$

$$= \sum_{k=1}^{n} \sum_{\substack{l=1 \\ l\neq k}}^{n} \left( \frac{2N_{i_{kr}}N_{i_{ls}}}{N_i^2} H_{i_{kr}i_{ls},t} + \frac{2N_{i_{ks}}N_{i_{lr}}}{N_i^2} H_{i_{kr}i_{ls},t} \right)$$

$$= \sum_{k=1}^{n} \sum_{\substack{l=1 \\ l\neq k}}^{n} \left( \frac{2f_{ik}t_{il}N_kN_l}{N_i^2} H_{i_{kr}i_{ls},t} + \frac{2t_{ik}f_{il}N_kN_l}{N_i^2} H_{i_{ks}i_{lr},t} \right) \tag{S24}$$

$$H_{ii,t}|_{2\cap B\cap\gamma} = \sum_{k=1}^{n} \sum_{\substack{l=1 \\ l\neq k}}^{n} \left( \frac{N_{i_{kr}}N_{i_{lr}}}{N_i^2} H_{i_{kr}j_{lr},t} + \frac{N_{i_{lr}}N_{i_{kr}}}{N_i^2} H_{i_{lr}j_{kr},t} \right)$$

$$= \sum_{k=1}^{n} \sum_{\substack{l=1 \\ l\neq k}}^{n} \left( \frac{2N_{i_{lr}}N_{i_{kr}}}{N_i^2} H_{i_{lr}j_{kr},t} \right)$$

$$= \sum_{k=1}^{n} \sum_{\substack{l=1 \\ l\neq k}}^{n} \left( \frac{2f_{ik}f_{il}N_kN_l}{N_i^2} H_{i_{lr}j_{kr},t} \right) \tag{S25}$$

As in the case of $H_{i_{ms}i_{ms},t}$, two genes are sampled from sub-stage $i_{ms}$ with replacement. Because all genes in sub-stage $i_{ms}$ were transferred by survival from stage $m$, sub-stage $i_{ms}$ consist of genes that were randomly sampled $2t_{im}N_m$ times 'without' replacement from stage $m$. Therefore, the probability of sampling two genes that are non-identical-by-descent without replacement should remain the same between stage $m$ in year $t-1$ and sub-stage $i_{ms}$ in year $t$ (i.e., $H'_{mm,t-1} = H'_{i_{ms}i_{ms},t}$). As with equation S5 , $H_{i_{ms}i_{ms},t}$ is formulated as follows.

$$H_{i_{ms}i_{ms},t} = \frac{1}{2t_{im}N_m} \times 0 + \left(1 - \frac{1}{2t_{im}N_m}\right) \times H'_{i_{ms}i_{ms},t}, \tag{S26}$$

From equations S5 and S26 ,

$$\begin{aligned} H'_{i_{ms}i_{ms},t} &= H'_{mm,t-1} \\ H_{i_{ms}i_{ms},t} &= \frac{1 - 1/(2t_{im}N_m)}{1 - 1/(2N_m)} H_{mm,t-1}. \end{aligned} \tag{S27}$$

It should be noted that $H_{i_{ms}i_{ms},t}$ should not be equal to $H_m m, t-1$. Two genes are always sampled from a common subset of stage $m$ (i.e, sub-stage $i_{ms}$), which means that two genes are not sampled from separate and independent surrogates of stage $m$ of the previous year. Therefore, sampling two genes from sub-stage $i_{ms}$ with replacement is not equivalent to that from stage $m$ with replacement. Unlike $H_{i_{ms}i_{ms},t}$, $H_{i_{ms}i_{mr},t}$ should be equal to $H_{mm,t-1}$, because sub-stage $i_m s$ and $i_m r$, from which two genes are sampled, are independently formed from stage $m$.

$$H_{i_{ms}i_{mr},t} = H_{mm,t-1} \tag{S28}$$

In the case of $H_{i_{mr}i_{mr},t}$, the sources of two genes sampled are the same (i.e., sub-stage $i_{mr}$) and thus are not independent surrogates of stage $m$ of the previous year, as with the case of $H_{i_{ms}i_{ms},t}$. Sampling the same gene twice occurs with the probability of $\frac{1}{2f_{im}N_m}$, which makes no contribution to $H_{i_{mr}i_{mr},t}$. In the remaining conditions where the two genes are sampled without replacement, the two genes are not identical-by-descent with a chance of $H_{mm,t-1}$ because sub-stage $i_{mr}$ were formed by reproduction, or sampling with replacement. Therefore, $H_{mm,t-1}$ is discounted by the fraction of $\frac{1}{2f_{im}N_m}$.

$$H_{i_{mr}i_{mr}} = \left(1 - \frac{1}{2f_{im}N_m}\right) H_{mm,t-1} \tag{S29}$$

In the case of $H_{i_{ks}i_{ls},t}$, $H_{i_{ks}i_{lr},t}$, $H_{i_{kr}i_{ls},t}$, and $H_{i_{kr}i_{lr},t}$, two genes are sampled from independent subset or copy of stage $k$ and $l$ of the previous year. Therefore,

$$H_{i_{ks}i_{ls},t} = H_{i_{ks}i_{lr},t} = H_{i_{kr}i_{ls},t} = H_{i_{kr}i_{lr},t} = H_{kl,t-1} \tag{S30}$$

Substituting equations S27 -S30 to equations S20 -S25 ,

$$H_{ii,t}|_{2\cap A\cap\alpha} = \sum_{m=1}^{n} \left\{ \left(\frac{t_{im}N_m}{N_i}\right)^2 \times \frac{1 - 1/(2t_{im}N_m)}{1 - 1/(2N_m)} H_{mm,t-1} \right\}$$

$$
\begin{aligned}
H_{ii,t}|_{2\cap A\cap\beta} &= \sum_{m=1}^{n}\left(\frac{2t_{im}f_{im}N_m^2}{N_i^2}\times H_{mm,t-1}\right)\\
H_{ii,t}|_{2\cap A\cap\gamma} &= \sum_{m=1}^{n}\left\{\left(\frac{f_{im}N_m}{N_i}\right)^2\times\left(1-\frac{1}{2f_{im}N_m}\right)H_{mm,t-1}\right\}\\
H_{ii,t}|_{2\cap B\cap\alpha} &= \sum_{k=1}^{n}\sum_{\substack{l=1\\l\neq k}}^{n}\left(\frac{2t_{ik}t_{il}N_kN_l}{N_i^2}\times H_{kl,t-1}\right)\\
H_{ii,t}|_{2\cap B\cap\beta} &= \sum_{k=1}^{n}\sum_{\substack{l=1\\l\neq k}}^{n}\left\{\frac{2(t_{ik}f_{il}+f_{ik}t_{il})N_kN_l}{N_i^2}\times H_{kl,t-1}\right\}\\
H_{ii,t}|_{2\cap B\cap\gamma} &= \sum_{k=1}^{n}\sum_{\substack{l=1\\l\neq k}}^{n}\left(\frac{2f_{ik}f_{il}N_kN_l}{N_i^2}\times H_{kl,t-1}\right).
\end{aligned}
\tag{S31}
$$

Substituting equations S31 to S19 , we can formulate $H_{ii,t}$.

$$
\begin{aligned}
H_{ii,t} = &\sum_{m=1}^{n}\left\{\left(\frac{t_{im}N_m}{N_i}\right)^2\frac{1-1/(2t_{im}N_m)}{1-1/(2N_m)}+\frac{2t_{im}f_{im}N_m^2}{N_i^2}\right.\\
&\left.+\left(\frac{f_{im}N_m}{N_i}\right)^2\left(1-\frac{1}{2f_{im}N_m}\right)\right\}H_{mm,t-1}\\
&+\sum_{k=1}^{n}\sum_{\substack{l=1\\l\neq k}}^{n}\frac{2(t_{ik}t_{il}+t_{ik}f_{il}+f_{ik}t_{il}+f_{ik}f_{il})N_kN_l}{N_i^2}H_{kl,t-1}\\
= &\sum_{m=1}^{n}\left\{\left(\frac{t_{im}N_m}{N_i}\right)^2\frac{1-1/(2t_{im}N_m)}{1-1/(2N_m)}+\frac{2t_{im}f_{im}N_m^2}{N_i^2}\right.\\
&\left.+\left(\frac{f_{im}N_m}{N_i}\right)^2\left(1-\frac{1}{2f_{im}N_m}\right)\right\}H_{mm,t-1}\\
&+\sum_{k=1}^{n}\sum_{\substack{l=1\\l\neq k}}^{n}\frac{2a_{ik}a_{il}N_kN_l}{N_i^2}H_{kl,t-1}.
\end{aligned}
\tag{S32}
$$

##### 1.3 Definition of generation time $T$

We use generation time $T$ to formulate effective population size $N_e$. Here, we explain the definition of generation time.

Firstly, we decompose the population projection matrix into two: $\boldsymbol{U}$ matrix, which is made up of $t_{ij}$ and describes the survival process, and $\boldsymbol{F}$ matrix, which is made up of stage-specific fecundity $f_{ij}$. In the case of the two-stage model,

$$\begin{pmatrix} t_{11} & t_{12}+f_{12} \\ t_{21} & t_{22} \end{pmatrix} = \begin{pmatrix} t_{11} & t_{12} \\ t_{21} & t_{22} \end{pmatrix} + \begin{pmatrix} 0 & f_{12} \\ 0 & 0 \end{pmatrix} = \boldsymbol{U} + \boldsymbol{F}. \tag{S33}$$

In the case of the three-stage model,

$$\begin{pmatrix} t_{11} & 0 & f_{13} \\ t_{21} & t_{22} & t_{23} \\ 0 & t_{32} & t_{33} \end{pmatrix} = \begin{pmatrix} t_{11} & 0 & 0 \\ t_{21} & t_{22} & t_{23} \\ 0 & t_{32} & t_{33} \end{pmatrix} + \begin{pmatrix} 0 & 0 & f_{13} \\ 0 & 0 & 0 \\ 0 & 0 & 0 \end{pmatrix} = \boldsymbol{U} + \boldsymbol{F}. \tag{S34}$$

By multiplying $\boldsymbol{U}$ matrix $x$ times, we can obtain transition probabilities per $x$ years. In the case of the two-stage model,

$$\boldsymbol{U}^x = \begin{pmatrix} t_{11} & t_{12} \\ t_{21} & t_{22} \end{pmatrix}^x = \begin{pmatrix} \tilde{u}_{11} & \tilde{u}_{12} \\ \tilde{u}_{21} & \tilde{u}_{22} \end{pmatrix}. \tag{S35}$$

Here, $\tilde{u}_{11}$ and $\tilde{u}_{21}$ are the probabilities that an individual in stage 1 remain in stage 1, or move to stage 2, after $x$ years, respectively. Now, we can formulate age-specific survival rate $l_x$, which denotes the probability of a newborn individual to survive until age $x$, and age-specific fecundity $m_x$, which is a expected number of newborns that an individual of age $x$ can make.

$$l_x = \tilde{u}_{11} + \tilde{u}_{21} \tag{S36}$$

$$m_x = 0 \times \frac{\tilde{u}_{11}}{\tilde{u}_{11} + \tilde{u}_{21}} + f_{12} \times \frac{\tilde{u}_{21}}{\tilde{u}_{11} + \tilde{u}_{21}}. \tag{S37}$$

In the case of the three-stage,

$$\boldsymbol{U}^x = \begin{pmatrix} t_{11} & 0 & 0 \\ t_{21} & t_{22} & t_{23} \\ 0 & t_{32} & t_{33} \end{pmatrix}^x = \begin{pmatrix} \tilde{u}_{11} & \tilde{u}_{12} & \tilde{u}_{13} \\ \tilde{u}_{21} & \tilde{u}_{22} & \tilde{u}_{23} \\ \tilde{u}_{31} & \tilde{u}_{32} & \tilde{u}_{33} \end{pmatrix} \tag{S38}$$

$$l_x = \tilde{u}_{11} + \tilde{u}_{21} + \tilde{u}_{31} \tag{S39}$$

$$m_x = f_{13} \times \frac{\tilde{u}_{31}}{\tilde{u}_{11} + \tilde{u}_{21} + \tilde{u}_{31}}. \tag{S40}$$

Then, we formulate generation time ($T$) as the expected age of a parent of a cohort.

$$T = \frac{\sum_{x=1}^{x_{max}} x l_x m_x}{\sum_{x=1}^{x_{max}} l_x m_x}, \tag{S41}$$

where $x_{max}$ is the maximum age defined as the age at which either of the two criteria (quoted from Waples et al., (2013)) is satisfied.

1. oldest age for which $l_x$ was $\geq 1$ % of the value at age at maturity ($L_\alpha$)
2. oldest age for which the product $l_x v_x$ was $\geq 1$ % of the maximum $l_x v_x$ for any age, where $v_x$ is the reproductive value of an individual of age $x$

Equation S41 is exactly the mean age of net fecundity in the cohort (Carey & Roach, 2020).

#### 2 Elements of matrix $M_2$ and $M_3$

Here, we show the elements of matrix $\boldsymbol{M_2}$ and $\boldsymbol{M_3}$. These two matrices correspond to the matrix $\boldsymbol{M}$ in equation 10 of the main text for the two- and three-stage model (Figure 2 of the main text), respectively.

### 2.1 $M_2$

$$\boldsymbol{h_t} = \begin{pmatrix} H_{11,t} \\ H_{22,t} \\ H_{12,t} \end{pmatrix} = \boldsymbol{M_2} \begin{pmatrix} H_{11,t-1} \\ H_{22,t-1} \\ H_{12,t-1} \end{pmatrix}$$

$$= \begin{pmatrix} m_{11} & m_{12} & m_{13} \\ m_{21} & m_{22} & m_{23} \\ m_{31} & m_{32} & m_{33} \end{pmatrix} \begin{pmatrix} H_{11,t-1} \\ H_{22,t-1} \\ H_{12,t-1} \end{pmatrix} \tag{S42}$$

where

$$m_{11} = t_{11}^2 \frac{1 - 1/(2t_{11}N_1)}{1 - 1/(2N_1)} \tag{S43}$$

$$m_{21} = \left(\frac{t_{21}N_1}{N_2}\right)^2 \frac{1 - 1/(2t_{21}N_1)}{1 - 1/(2N_1)} \tag{S44}$$

$$m_{31} = \frac{N_1}{N_2} \left(\frac{t_{11}t_{21}}{1 - 1/(2N_1)}\right) \tag{S45}$$

$$m_{12} = \left(\frac{t_{12}N_2}{N_1}\right)^2 \frac{1 - 1/(2t_{12}N_2)}{1 - 1/(2N_2)} + \frac{2t_{12}f_{12}N_2^2}{N_1^2} + \left(\frac{f_{12}N_2}{N_1}\right)^2 \left(1 - \frac{1}{2f_{12}N_2}\right) \tag{S46}$$

$$m_{22} = t_{22}^2 \frac{1 - 1/(2t_{22}N_2)}{1 - 1/(2N_2)} \tag{S47}$$

$$m_{32} = \frac{N_2}{N_1} \left(\frac{t_{12}t_{22}}{1 - 1/(2N_2)} + f_{12}t_{22}\right) \tag{S48}$$

$$m_{13} = \frac{2t_{11}a_{12}N_2}{N_1} \tag{S49}$$

$$m_{23} = \frac{2t_{21}t_{22}N_1}{N_2} \tag{S50}$$

$$m_{33} = t_{11}t_{22} + a_{12}t_{21} \tag{S51}$$

#### 2.2 $\boldsymbol{M_3}$

$$\boldsymbol{h_t} = \begin{pmatrix} H_{11,t} \\ H_{22,t} \\ H_{33,t} \\ H_{12,t} \\ H_{23,t} \\ H_{13,t} \end{pmatrix} = \boldsymbol{M_3} \begin{pmatrix} H_{11,t-1} \\ H_{22,t-1} \\ H_{33,t-1} \\ H_{12,t-1} \\ H_{23,t-1} \\ H_{13,t-1} \end{pmatrix}$$

$$= \begin{pmatrix} m_{11} & m_{12} & m_{13} & m_{14} & m_{15} & m_{16} \\ m_{21} & m_{22} & m_{23} & m_{24} & m_{25} & m_{26} \\ m_{31} & m_{32} & m_{33} & m_{34} & m_{35} & m_{36} \\ m_{41} & m_{32} & m_{43} & m_{44} & m_{45} & m_{46} \\ m_{51} & m_{42} & m_{53} & m_{54} & m_{55} & m_{56} \\ m_{61} & m_{52} & m_{63} & m_{64} & m_{65} & m_{66} \end{pmatrix} \begin{pmatrix} H_{11,t-1} \\ H_{22,t-1} \\ H_{33,t-1} \\ H_{12,t-1} \\ H_{23,t-1} \\ H_{13,t-1} \end{pmatrix} \tag{S52}$$

where

$$m_{11} = t_{11}^2 \frac{1 - 1/(2t_{11}N_1)}{1 - 1/(2N_1)} \tag{S53}$$

$$m_{21} = \left(\frac{t_{21}N_1}{N_2}\right)^2 \frac{1 - 1/(2t_{21}N_1)}{1 - 1/(2N_1)} \tag{S54}$$

$$m_{31} = 0 \tag{S55}$$

$$m_{41} = \frac{N_1}{N_2} \frac{t_{11} t_{21}}{1 - 1/(2N_1)} \tag{S56}$$

$$m_{51} = 0 \tag{S57}$$

$$m_{61} = 0 \tag{S58}$$

$$m_{12} = 0 \tag{S59}$$

$$m_{22} = t_{22}^2 \frac{1 - 1/(2t_{22}N_2)}{1 - 1/(2N_2)} \tag{S60}$$

$$m_{32} = \left(\frac{t_{32}N_2}{N_3}\right)^2 \frac{1 - 1/(2t_{32}N_2)}{1 - 1/(2N_2)} \tag{S61}$$

$$m_{42} = 0 \tag{S62}$$

$$m_{52} = \frac{N_2}{N_3} \frac{t_{22} t_{32}}{1 - 1/(2N_2)} \tag{S63}$$

$$m_{62} = 0 \tag{S64}$$

$$m_{13} = \left(\frac{f_{13}N_3}{N_1}\right)^2 \left(1 - \frac{1}{2f_{13}N_3}\right) \tag{S65}$$

$$m_{23} = \left(\frac{t_{23}N_3}{N_2}\right)^2 \frac{1 - 1/(2t_{23}N_3)}{1 - 1/(2N_3)} \tag{S66}$$

$$m_{33} = t_{33}^2 \frac{1 - 1/(2t_{33}N_3)}{1 - 1/(2N_3)} \tag{S67}$$

$$m_{43} = \frac{f_{13} t_{23} N_3^2}{N_1 N_2} \tag{S68}$$

$$m_{53} = \frac{N_3}{N_2} \frac{t_{23} t_{33}}{1 - 1/(2N_3)} \tag{S69}$$

$$m_{63} = \frac{f_{13} t_{33} N_3}{N_1} \tag{S70}$$

$$m_{14} = 0 \tag{S71}$$

$$m_{24} = \frac{2 t_{21} t_{22} N_1}{N_2} \tag{S72}$$

$$m_{34} = 0 \tag{S73}$$

$$m_{44} = t_{11} t_{22} \tag{S74}$$

$$m_{54} = \frac{t_{21} t_{32} N_1}{N_3} \tag{S75}$$

$$m_{64} = \frac{t_{11} t_{32} N_2}{N_3} \tag{S76}$$

$$m_{15} = 0 \tag{S77}$$

$$m_{25} = \frac{2 t_{22} t_{23} N_3}{N_2} \tag{S78}$$

$$m_{35} = \frac{t_{32} t_{33} N_2}{N_3} \tag{S79}$$

$$m_{45} = \frac{f_{13} t_{22} N_3}{N_1} \tag{S80}$$

$$m_{55} = t_{22}t_{33} \tag{S81}$$

$$m_{65} = \frac{t_{32}f_{13}N_2}{N_1} \tag{S82}$$

$$m_{16} = \frac{2t_{11}f_{13}N_3}{N_1} \tag{S83}$$

$$m_{26} = \frac{2t_{21}t_{23}N_1N_3}{N_2^2} \tag{S84}$$

$$m_{36} = 0 \tag{S85}$$

$$m_{46} = \frac{f_{13}t_{21}N_3}{N_2} \tag{S86}$$

$$m_{56} = \frac{t_{21}t_{33}N_1}{N_2} \tag{S87}$$

$$m_{66} = t_{11}t_{33} \tag{S88}$$

#### 3 How to determine parameter values

We randomly produced 500 parameter sets for each of the two- and the three-stage model for simulation and model analysis. Here, we explain how we determined the values of each parameter (demographic rates and the number of individuals in each stage). We also showed an example case in Figure S1.

**Step 1** We draw four random numbers from the uniform distribution $U(0, 1)$ for the two-stage model. In the case of the three-stage model, six random numbers are drawn from the same uniform distribution.

**Step 2** We rearrange the random numbers in an increasing order.

**Step 3** We multiply the random numbers by 100, and round them off to be integers. Moreover, we add 0 and 100 to the sequences.

**Step 4** We take the difference between the neighboring numbers: we subtract each number from its next smaller one. As a result, five and seven numbers are generated for the two-stage and the three-stage models, respectively.

**Step 5** Each number is assigned to one of the demographic processes (i.e., growth, stasis, retrogression, and reproduction) of each stage. As for the two-stage model, the first to fifth numbers are assigned to (1) stasis at juvenile, (2) growth from juvenile to adult, (3) retrogression from adult to juvenile, (4) stasis at adult, and (5) reproduction, respectively. In the case of the three-stage model, seven numbers are sequentially assigned to (1) stasis at seed, (2) growth from seed to juvenile, (3) stasis at juvenile, (4) growth from juvenile to adult, (5) retrogression from adult to juvenile, (6) stasis at adult, and (7) reproduction.

**Step 6** We calculate the number of individuals of each stage as the sum of flows coming into each stage.

**Step 7** We calculate demographic rates by dividing the number of individuals of corresponding flows by that of the stages from which the flows come out.

**Step 8** We assess if the parameter values calculated in step 7 completely satisfy the following three criteria. If they do, the values are added to the parameter sets. If not, the values are discarded and we restart the procedures from Step 1.

1. Growth probability and fecundity should be greater than 0, otherwise the life cycle would be broken off.
2. Survival probability of each stage (i.e., sum of growth, stasis, and retrogression probabilities of each stage) is within the range of [0, 1] to be a probability.
3. At least one stasis probability is not zero, otherwise genes in different stages do not mix with one another and are segregated eternally.

**Step 9** We repeated Step 1 to 8 until the number of parameter sets reached 500.

The resultant parameter sets range the parameter space widely both for the two-stage and for the three-stage models, indicating that our model is examined for a variety of demographic strategies (Figure S2).

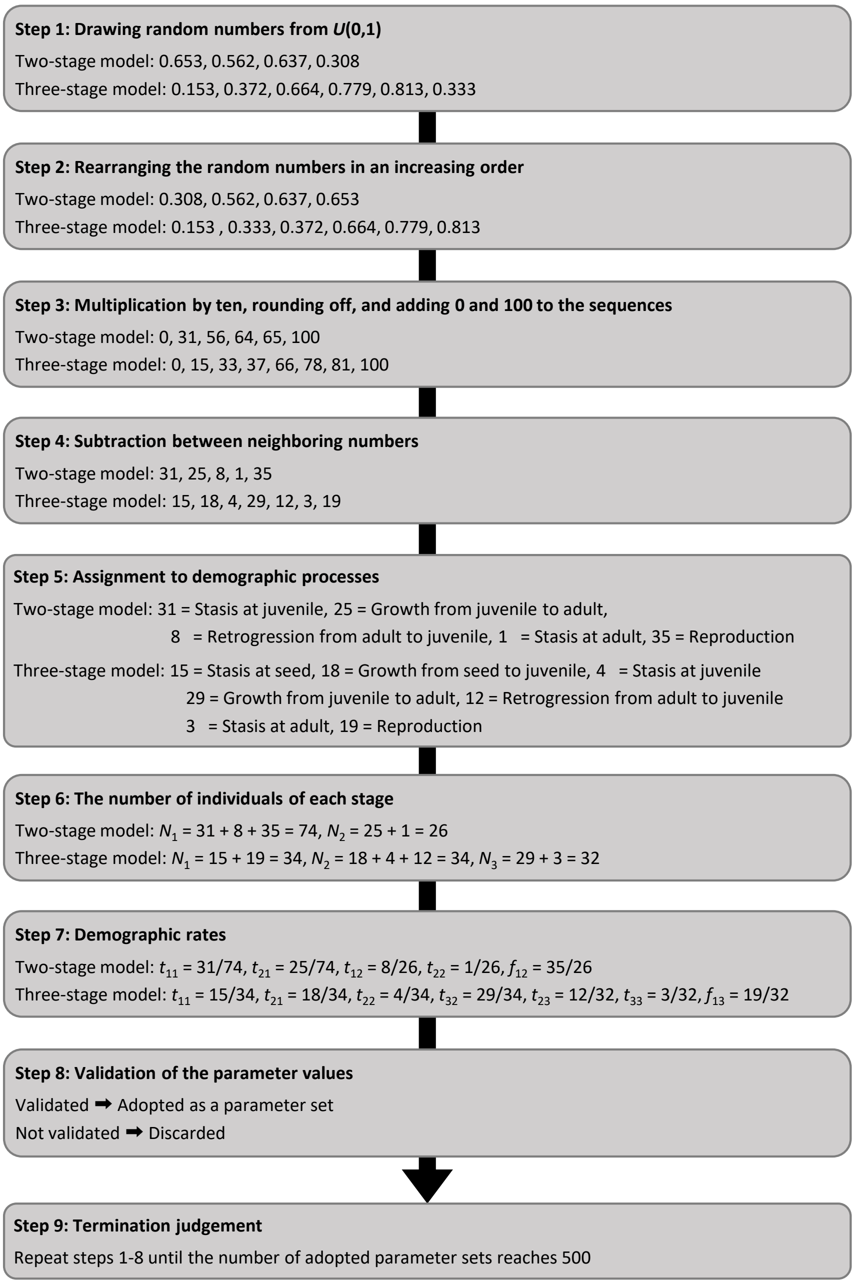

Figure S 1: A graphical view of steps 1 to 9 with an example for the two- and the three-stage models

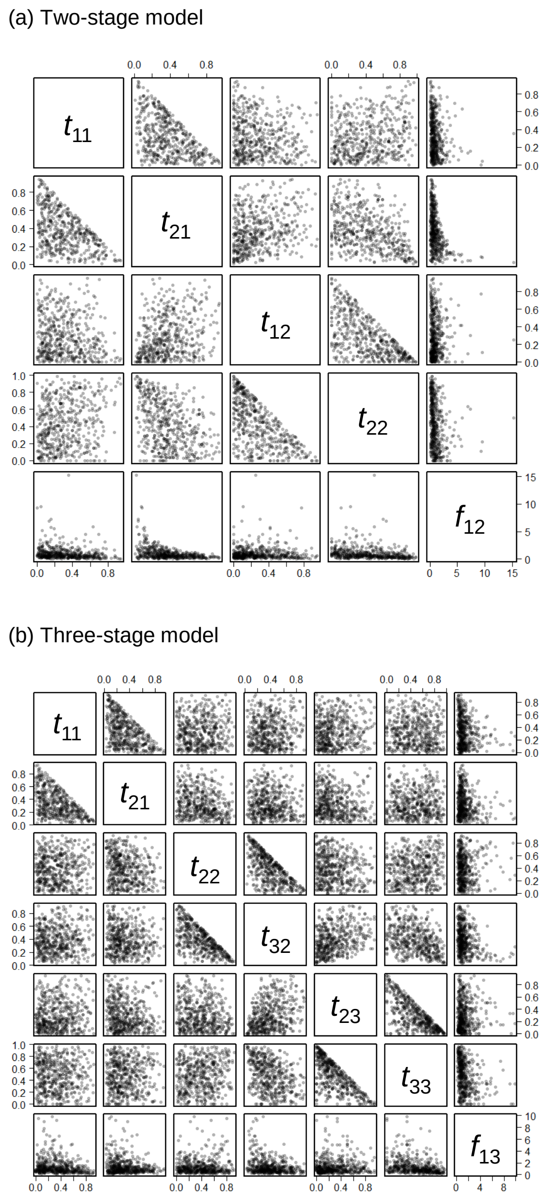

Figure S 2: 500 parameter sets used in simulation and model analysis. One dot corresponds to one parameter set. (a) Two-stage model, (b) three-stage model. There are some parameter pairs where the dots occupy only the lower-left triangle (e.g., $t_{11}$ and $t_{21}$ in (a); $t_{22}$ and $t_{32}$ in (b)). This is because of the second criterion in step 8, that is, the sum should not exceed 1

### 4 Additional results

#### 4.1 Validation of our model

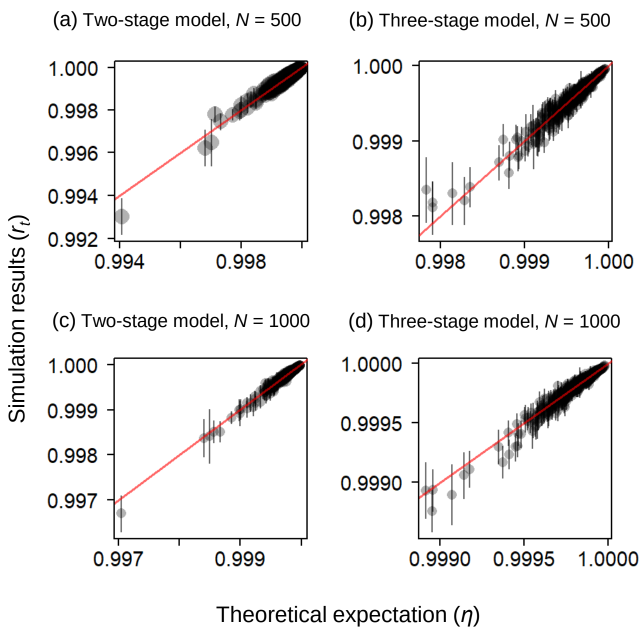

Figure S 3: We compared the rate of change in expected heterozygosity between theoretical expectation ($\eta$) and simulation results ($r_t$) when $N$ =500 and 1000 for both the two- and the three-stage model. (a) Two-stage model, $N$ = 500, (b) three-stage model, $N$ = 500, (c) two-stage model, $N$ = 1000, (d) three-stage model, $N$ = 1000. Vertical bars represent standard error of $r_t$. Red lines indicate $r_t = \eta$

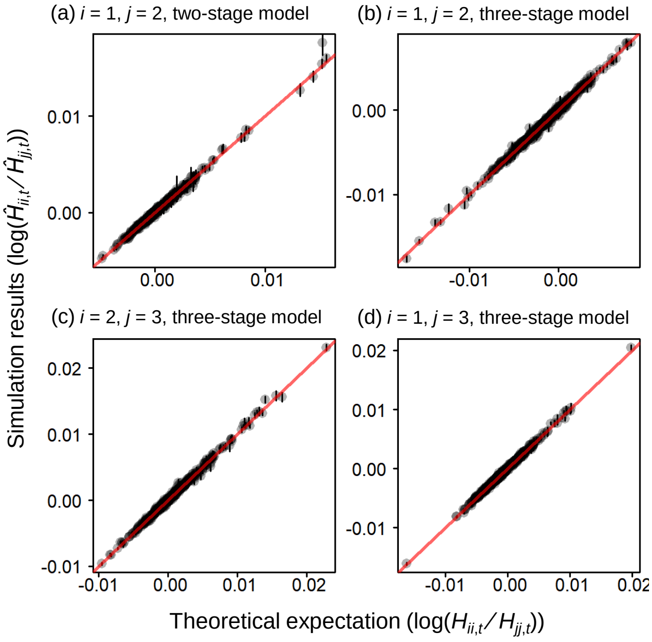

Figure S 4: Comparison of demographic genetic structure between theoretical expectation ($\log(H_{ii,t}/H_{jj,t})$) and simulation results ($\log(\hat{H}_{ii,t}/\hat{H}_{jj,t})$) when $N$ =500 for both the two- and the three-stage model. (a) $i = 1$, $j = 2$, two-stage model, (b)$i = 1$, $j = 2$, three-stage model, (c) $i = 2$, $j = 3$, three-stage model, (d) $i = 1$, $j = 3$, three-stage model. Vertical bars represent standard error of $r_t$. Red lines indicate $\log(H_{ii,t}/H_{jj,t})$=$\log(\hat{H}_{ii,t}/\hat{H}_{jj,t})$

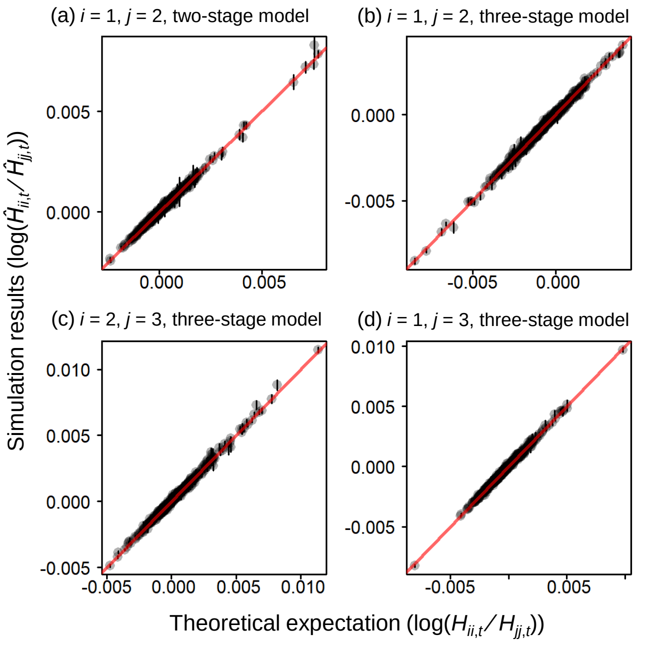

Figure S 5: Comparison of demographic genetic structure between theoretical expectation ($\log(H_{ii,t}/H_{jj,t})$) and simulation results ($\log(\hat{H}_{ii,t}/\hat{H}_{jj,t})$) when $N$ =1000 for both the two- and the three-stage model. (a) $i = 1$, $j = 2$, two-stage model, (b)$i = 1$, $j = 2$, three-stage model, (c) $i = 2$, $j = 3$, three-stage model, (d) $i = 1$, $j = 3$, three-stage model. Vertical bars represent standard error of $r_t$. Red lines indicate $\log(H_{ii,t}/H_{jj,t})$=$\log(\hat{H}_{ii,t}/\hat{H}_{jj,t})$

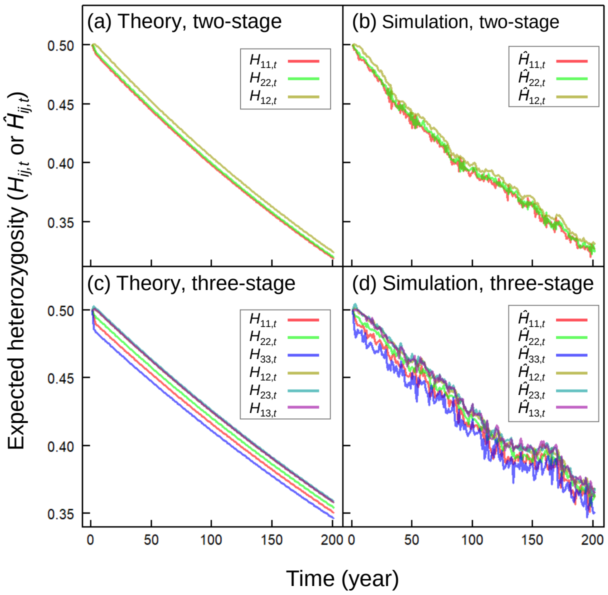

Figure S 6: Graphical comparison of tempotal dynamics of expected heterozygosity between theoretical expectation (i.e., dynamics of $H_{ij,t}$) and simulation results (i.e., dynamics of $\hat{H}_{ij,t}$) under a partocular parameter set in the two- and the three-stage model. Colors stand for combinations of $i$ and $j$. (a) Theoretical expectations and (b) simulation results of the two-stage model, (c) theoretical expectations and (d) simulation results of the three-stage model. Parameter set of the two-stage model is $t_{11} = 0.115, t_{21} = 0.750, t_{12} = 0.333, t_{22} = 0.188, f_{12} = 0.625, N_1 = 52, N_2 = 48$, while that of the three-stage model is $t_{11} = 0.476, t_{21} = 0.405, t_{22} = 0.568, t_{32} = 0.273, t_{23} = 0.143, t_{33} = 0.143, f_{13} = 1.57, N_1 = 42, N_2 = 44, N_3 = 14$. Each parameter set was randomly picked from the 500 parameter sets under $N = 100$ as an example case

#### 4.2 Comparison of demographic genetic structure with $N_e$ and $\eta$

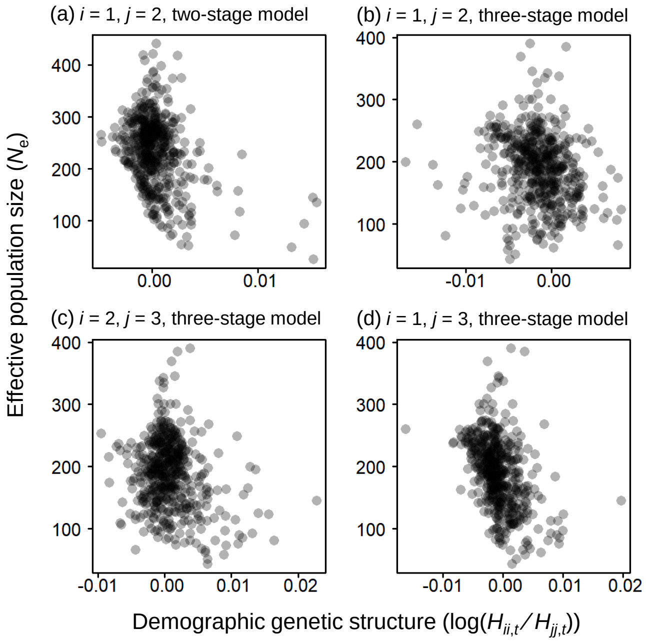

Figure S 7: Comparison of demographic genetic structure ($\log(H_{ii,t}/H_{jj,t})$) with effective population size ($N_e$) when $N = 500$. (a) $i = 1$ and $j = 2$ of the two-stage model, (b) $i = 1$ and $j = 2$, (c) $i = 2$ and $j = 3$, (d) $i = 1$ and $j = 3$ of the three-stage model

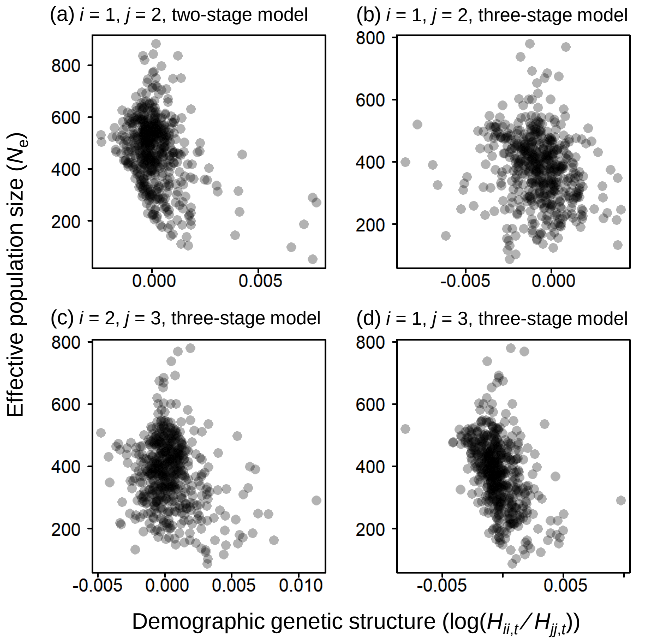

Figure S 8: Comparison of demographic genetic structure ($\log(H_{ii,t}/H_{jj,t})$) with effective population size ($N_e$) when $N = 1000$. (a) $i = 1$ and $j = 2$ of the two-stage model, (b) $i = 1$ and $j = 2$, (c) $i = 2$ and $j = 3$, (d) $i = 1$ and $j = 3$ of the three-stage model

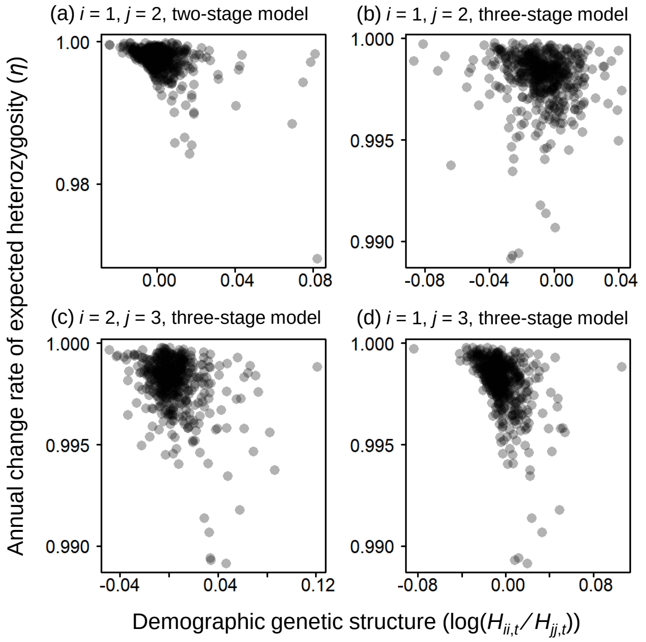

Figure S 9: Comparison of demographic genetic structure ($\log(H_{ii,t}/H_{jj,t})$) with the anual change rate of expected heterozygosity ($\eta$) when $N = 100$. (a) $i = 1$ and $j = 2$ of the two-stage model, (b) $i = 1$ and $j = 2$, (c) $i = 2$ and $j = 3$, (d) $i = 1$ and $j = 3$ of the three-stage model

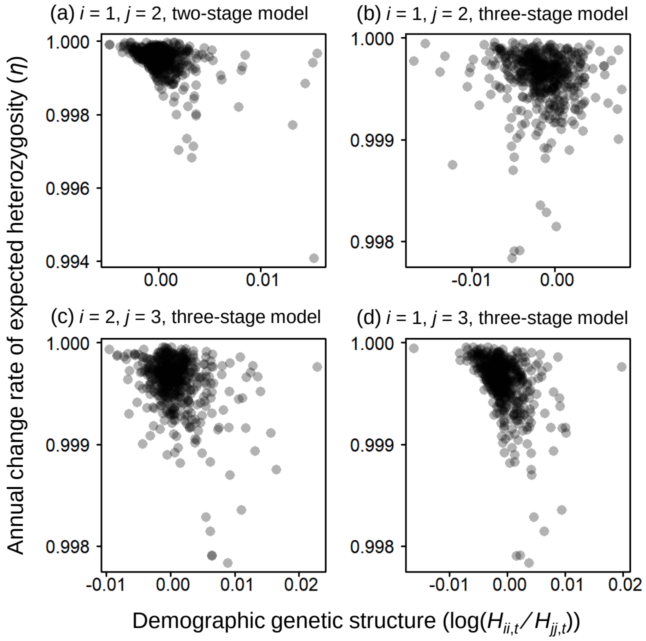

Figure S 10: Comparison of demographic genetic structure ($\log(H_{ii,t}/H_{jj,t})$) with the anual change rate of expected heterozygosity ($\eta$) when $N = 500$. (a) $i = 1$ and $j = 2$ of the two-stage model, (b) $i = 1$ and $j = 2$, (c) $i = 2$ and $j = 3$, (d) $i = 1$ and $j = 3$ of the three-stage model

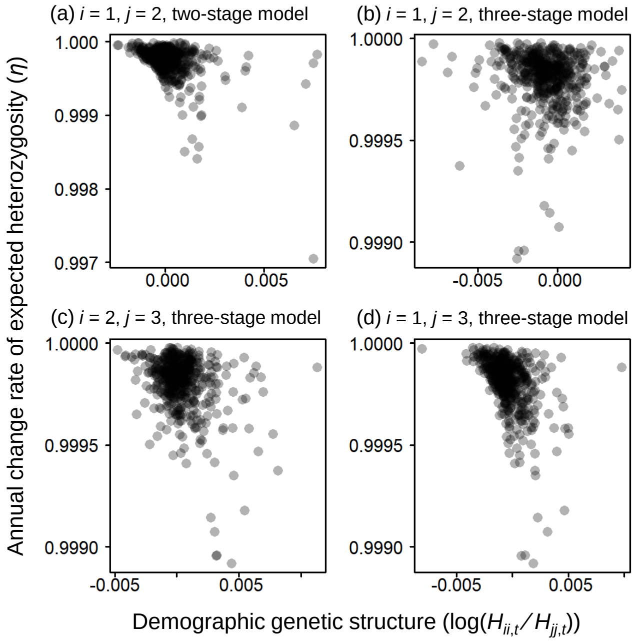

Figure S 11: Comparison of demographic genetic structure ($\log(H_{ii,t}/H_{jj,t})$) with the anual change rate of expected heterozygosity ($\eta$) when $N = 1000$. (a) $i = 1$ and $j = 2$ of the two-stage model, (b) $i = 1$ and $j = 2$, (c) $i = 2$ and $j = 3$, (d) $i = 1$ and $j = 3$ of the three-stage model

#### 4.3 Dependence of $N_e$ and $\eta$ to $N$

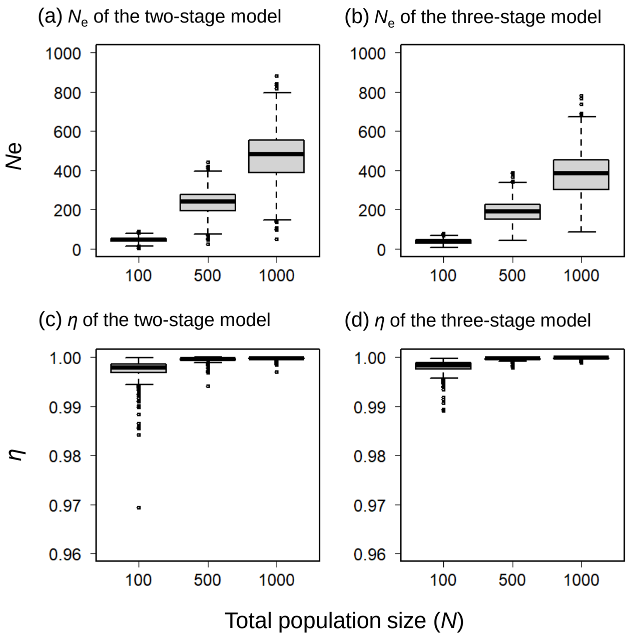

Figure S 12: Effective population size ($N_e$) and the annual change rate of expected heterozygosity ($\eta$) of the 500 parameter sets for each of $N = 100, 500$, and 1000. (a) $N_e$ of the two-stage model, (b) $N_e$ of the three-stage model, (c) $\eta$ of the two-stage model, (d) $\eta$ of the three-stage model
